# Diversity, distribution, and expression of opsin genes in freshwater lakes

**DOI:** 10.1101/2022.08.01.502354

**Authors:** Shaomei He, Alexandra M. Linz, Sarah L.R. Stevens, Patricia Q. Tran, Francisco Moya-Flores, Ben O. Oyserman, Jeffrey R. Dwulit-Smith, Katrina T. Forest, Katherine D. McMahon

## Abstract

Microbial rhodopsins are widely distributed in aquatic environments and may significantly contribute to phototrophy and energy budgets in global oceans. However, the study of freshwater rhodopsins has been largely limited. Here, we explored the diversity, ecological distribution, and expression of opsin genes that encode the apoproteins of Type I rhodopsins in humic and clearwater lakes with contrasting physicochemical and optical characteristics. Using metagenomes and metagenome-assembled genomes, we recovered opsin genes from a wide range of taxa, mostly predicted to encode green light-absorbing proton pumps. Viral opsin and novel bacterial opsin clades were recovered. Opsin genes occurred more frequently in taxa from clearwater than from humic water, and opsins in some taxa have non-typical ion-pumping motifs that might be associated with physicochemical conditions of these two freshwater types. Analyses of the surface layer of 33 freshwater systems revealed an inverse correlation between opsin gene abundance and lake dissolved organic carbon (DOC). In humic water with high terrestrial DOC and light-absorbing humic substances, opsin gene abundance was low and dramatically declined within the first few meters, whereas the abundance remained relatively high along the bulk water column in clearwater lakes with low DOC, suggesting opsin gene distribution is influenced by lake optical properties and DOC. Gene expression analysis confirmed the significance of rhodopsin-based phototrophy in clearwater lakes and revealed different diel expressional patterns among major phyla. Overall, our analyses revealed freshwater opsin diversity, distribution and expression patterns, and suggested the significance of rhodopsin-based phototrophy in freshwater energy budgets, especially in clearwater lakes.

## INTRODUCTION

Microbial rhodopsin is a retinal-activated photosystem. In marine environments, microbial rhodopsins are found in many diverse prokaryotes and exhibit a rich genetic and functional diversity (McCarren & DeLong 2007; Olson *et al*. 2018; Rozenberg *et al*. 2021; Rusch *et al*. 2007). As microbial rhodopsins often occur in more than half of the cells in a wide range of marine samples (Finkel *et al*. 2013; Maresca *et al*. 2018; Olson *et al*. 2018; Sieradzki *et al*. 2018), rhodopsin-based phototrophy has been suggested to be a prevalent form of phototrophy and may significantly influence energy budgets in the global oceans (Finkel *et al*. 2013; Gomez-Consarnau *et al*. 2019), although whether the harvested sunlight is used for growth, survival or other physiological and ecological benefits may differ among bacterial species (Pinhassi *et al*. 2016). Contrasting to the rich knowledge regarding marine microbial rhodopsins, the diversity, ecological distribution, and significance of rhodopsins in freshwater have been much less studied. Particularly, rhodopsins in humic aquatic environments (with high terrestrially-derived allochthonous organic carbon, dimly lit water columns, and often acidic pH) are largely unexplored.

Being a simple photosystem, a rhodopsin consists of an opsin, which is an integral membrane protein with seven transmembrane helices, and a covalently bound retinal as a light-absorbing chromophore. Microbial (also termed Type I, as compared to Type II in animals) rhodopsins play important roles in microbial physiology, with known functions including light-driven ion transport (e.g. proton, H^+^, chloride, Cl^-^, and sodium, Na^+^ pumps), cation channeling, and sensing that directs phototaxis (Rozenberg *et al*. 2021). More recently, a novel family of microbial rhodopsins distinct from Type I rhodopsins was discovered through functional metagenomics and named heliorhodopsin (HeR) (Pushkarev *et al*. 2018). As HeRs are only distantly related to Type I rhodopsins and their functions are largely unknown, in our study, we solely focused on Type I rhodopsins (hereinafter referred to as “rhodopsins” and genes encoding their apoproteins, opsins, were referred to as “opsin genes” for simplicity), although we note that HeRs have been found in diverse monoderm bacteria in freshwater environments (Chazan *et al*. 2022; Flores-Uribe *et al*. 2019).

The first microbial opsin genes in freshwater were discovered through metagenomic studies, where the predominant opsin genes were recovered from Actinobacteria and therefore named “actinorhodopsin” (ActR) (Sharma *et al*. 2009; Sharma *et al*. 2008). ActR is phylogenetically closer to xanthorhodopsins than to other rhodopsin groups (Vollmers *et al*. 2013) and functions as an outward H^+^ pump (Dwulit-Smith *et al*. 2018; Keffer *et al*. 2015). In addition to ActR, opsin genes were also recovered from some other phyla in freshwater lakes (Atamna-Ismaeel *et al*. 2008; Cabello-Yeves *et al*. 2017; Galachyants *et al*. 2021; Jaffe *et al*. 2022; Martinez-Garcia *et al*. 2012; Podowski *et al*. 2021; Sharma *et al*. 2009).

Rhodopsin has been shown to enhance bacterial growth and survival under carbon-limited conditions when exposed to light (Gómez-Consarnau *et al*. 2010; Gómez-Consarnau *et al*. 2007; Johnson *et al*. 2010; Steindler *et al*. 2011; Yoshizawa *et al*. 2014), suggesting the importance of light and carbon availability in the regulation of this photoreceptive protein. In freshwater, lake optical properties and organic carbon quality/quantity are often correlated, and the former can be greatly impacted by the latter (Maizel *et al*. 2017; Read & Rose 2013). Therefore, an interesting yet unexplored hypothesis is that the diversity and distribution of opsin genes in freshwater varies with different lake optical properties and organic carbon quality/quantity.

In this study, we surveyed two contrasting freshwaters, a eutrophic clearwater lake and a humic brown-water bog lake with distinct physicochemical characteristics. In particular, the humic bog has much higher concentrations of dissolved organic carbon (DOC), most of which is allochthonous (entering the aquatic system from its catchment). Such terrestrially-derived DOC comprises light-absorbing humic substances which stain the water brown, and thus the bog exhibits very different optical properties from the eutrophic lake, which is “clear” as opposed to “stained” (Maizel *et al*. 2017). To investigate the opsin gene diversity in these environments, we used metagenomic sequences and metagenome-assembled genomes (MAGs) obtained from the co-assembly of a large number of metagenomes collected from a time-series study of these two lakes (Linz *et al*. 2018), as well as the 12665 MAGs from the stratfreshDB which was generated from 41 lakes and ponds (Buck *et al*. 2021). To further study the relationship between opsin gene distribution and freshwater DOC and optical properties, we included metagenomes from a total of 33 freshwater lakes around the globe that span a wide range of DOC levels and clarity. We estimated the relative opsin gene abundance in the surface layer of these lakes, as well as depth-resolved opsin abundance profiles from a subset of these lakes. Lastly, we explored opsin gene expression and diel regulation in the epilimnion of two clearwater lakes and the humic bog. Overall, our analysis recovered a greater than previously appreciated diversity of freshwater opsin genes, revealed the importance of DOC and optical properties in opsin distribution, and suggested the significance of rhodopsin-based phototrophy in clearwater lakes.

## MATERIALS AND METHODS

### Lake Mendota and Trout Bog metagenomes

Two contrasting temperate lakes in Wisconsin, USA were studied in detail. Lake Mendota (43.10°N, 89.41°W) is a eutrophic clearwater lake with an average DOC of 5.0 mg/L. Trout Bog (46.01°N, 89.61°W) is a small humic lake, with an average DOC of 20.0 mg/L (data from NTL-LTER, averaged over the course of the study). More detailed lake physicochemical and limnological characteristics are listed in **Table S1**. For Lake Mendota, 0 to 12 m depth-integrated water samples (mostly consisting of the epilimnion layer) were collected at 94 time points from 2008 to 2012, and samples are referred to as “ME”. For Trout Bog, the integrated epilimnion layer and the integrated hypolimnion layer were collected at 45 time points respectively from 2007 to 2009, and samples were referred to as “TE” and “TH”, respectively.

Details of metagenome sequencing, assembly, and binning were previously described (Bendall *et al*. 2016; Linz *et al*. 2018). Briefly, shotgun metagenome libraries were constructed from each of the 184 DNA samples. Assembly was performed on individual metagenomes. In addition, three “combined” assemblies were generated by co-assembling reads from all metagenomes within the ME, TE, and TH groups, respectively. Binning was conducted on the three combined assemblies to obtain metagenome-assembled genomes (MAGs), and this resulted in 486 genome bins. By further fine-tuning to remove outlier contigs, and excluding bins with high contamination (>10%) or low completeness (<50%), a total of 202 high-quality MAGs (referred to henceforth as MAGs to distinguish them from the remaining bins which have lower qualities) were generated (Linz *et al*. 2018). Taxonomic assignment of MAGs and bins was conducted using PhyloSift based on 37 conserved phylogenetic marker genes (Darling *et al*. 2014). The 184 individual metagenome assemblies, the three combined assemblies, and recovered MAGs/bins were submitted to the DOE Joint Genome Institute’s Integrated Microbial Genome (IMG) database for gene prediction and function annotation (Markowitz *et al*. 2013).

### Opsin phylogenetic tree reconstruction and inference of taxonomy

Opsin sequences were identified by hmmsearch with model-specific trusted cut-offs (--cut_tc) for the hidden Markov model (HMM) of pfam01036 (http://pfam.xfam.org) implemented by the IMG annotation pipeline. Representative opsin reference sequences were chosen by clustering bacterial and archaeal opsin protein sequences from isolate genomes at IMG based on 90% amino acid sequence identity using the UCLUST algorithm provided by the USEARCH tool (v5.2.32) (Edgar 2010). Representative sequences from each cluster, opsins from previously sequenced freshwater single-amplified genomes (SAGs) and MAGs, and viral opsin sequences from the study by Yutin and Koonin (Yutin & Koonin 2012) were included in the phylogenetic analysis, together with the 317 opsin sequences longer than 200 amino acids (which is ∼80% of opsin full length) in our three combined assemblies (including sequences in MAGs/bins and the remaining un-binned). Amino acid sequences were aligned using MUSCLE (v3.8.31) (Edgar 2004), and the alignment was trimmed to exclude columns that contained gaps for more than 30% of the included sequences. Poorly aligned positions and divergent regions were further eliminated using Gblocks (v0.91b) (Castresana 2000). A maximum likelihood phylogenetic tree was constructed using PhyML 3.0 (Guindon *et al*. 2010), with the LG substitution model (Le & Gascuel 2008) and the gamma distribution parameter estimated by PhyML. Bootstrap values were calculated based on 100 replicates. After constructing this phylogenetic tree, sequences shorter than 200 amino acids in the three combined assemblies were added to the existing tree using PPlacer (<u>v1.1.alpha17</u>) (Matsen *et al*. 2010). The resulting phylogenetic tree was used for taxonomic classification of un-binned metagenome opsin sequences at the phylum level.

### Searching opsin genes in stratfreshDB MAGs

We first performed gene prediction using Prodigal (v2.6.3) on all stratfreshDB MAGs deposited at European Nucleotide Archive (Buck *et al*. 2021). We then performed hmmsearch (HMMER v3.3) with the model-specific trusted cutoff (--cut_tc) option for pfam01036 to identify opsin genes. Taxonomic information of these MAGs was directly from the classification with GTDB-Tk in the original study (Buck *et al*. 2021).

### Estimation of opsin gene relative abundance in ME, TE, and TH metagenomes

The abundance of an individual gene in an assembled metagenome was estimated using its average per-base coverage, which indicates how many times a base was sequenced and was determined by mapping metagenome reads to assembled contigs. This estimate is available in the IMG database as “read depth”, and we referred to it as “coverage-weighted abundance” in this study to reflect the abundance of the population that contributed to this gene in the metagenome. To compare opsin gene abundance among different metagenomes, the opsin gene relative abundance was estimated by the sum of the coverage-weighted abundance of all opsin genes in the metagenome, and further normalized (i.e. divided) by the average of the coverage-weighted abundance of three single-copy conserved housekeeping genes (represented by pfam00121, pfam00573, pfam00687, http://pfam.xfam.org) in the same metagenome. The three single-copy genes were chosen because they have comparable HMM lengths to opsin genes to minimize the bias in gene abundance estimates associated with gene lengths (He *et al*. 2015).

### Estimation of opsin gene abundance for additional lakes

To further study opsin gene distribution, we included metagenomes from 31 additional freshwater lakes for which organic carbon concentrations were available. Among these additional lakes, 18 are from the stratfreshDB deposited at European Nucleotide Archive (Buck *et al*. 2021) and 13 are from publicly available metagenomes at IMG, including two unpublished from our lab and 11 from published studies (Dam *et al*. 2020; Denef *et al*. 2016; Linz *et al*. 2020; Podowski *et al*. 2021; Tran *et al*. 2018; Tran *et al*. 2021) (**Tables S2 and S3**). Together with Mendota and Trout Bog, the 33 lakes span a wide range of DOC levels and exhibit different clarity (examples of clarity data are listed in **Table S1**). Some of these lakes have samples collected from different depths, allowing the study of opsin gene vertical distribution (**Table S4**). For metagenomes available at IMG, the relative abundance of opsin genes was estimated as described above. For metagenomes from the stratfreshDB, we performed gene prediction using Prodigal (v2.6.3) on the assemblies, and then hmmsearch (HMMER v3.3) with the model-specific trusted cutoff (--cut_tc) option for the HMM models of pfam01036, pfam00121, pfam00573, and pfam00687 to identify opsin and the three single-copy conserved genes respectively. To obtain gene coverage information, we first processed the raw reads to remove adapter sequences and low-quality bases using fastp (v0.23.1) with default settings. We then mapped the processed reads to corresponding metagenome assemblies using bowtie2 (v2.2.2) with default settings for gene coverage. Coverage-weighted abundance from these genes was used to estimate opsin gene abundance as described earlier. The in-house bioinformatics workflow that we used on the stratfreshDB metagenomes, when used on a subset of the included IMG metagenomes, yielded results very comparable to results obtained from the IMG pipeline. Therefore, no systemic bias was associated with different bioinformatic workflows.

### Analysis of opsin gene expression and diel trends

We used metatranscriptomes from the GEODES (Gene Expression in Oligotrophic, Dystrophic, and Eutrophic Systems) dataset (Linz *et al*. 2020) to study the expression of opsin genes. This dataset includes triplicate samples collected every four hours for two days from Lake Mendota and Sparking Lake, an oligotrophic clearwater lake in Northern Wisconsin (**Table S1**). Triplicate samples were similarly collected from Trout Bog but samples from the second day did not yield sufficient RNA for sequencing. Notably, an internal standard, an *in vitro* transcribed pFN18A of a known concentration was added to each sample after the bead beating step during the RNA extraction (Linz *et al*. 2020). This type of internal standard enables absolute quantification of transcript abundance as copies of transcripts per liter of aquatic sample and thus allows comparing transcript abundance across different samples (Gifford *et al*. 2011). As two metagenomes were collected along with the metatranscriptomes for each lake, we first clustered opsin sequences from the two replicate metagenomes with 97% nucleotide identity using the UCLUST algorithm provided by the USEARCH tool (v3.8.31) (Edgar 2010) to remove redundancy, followed by removing sequences shorter than 600 bp. The resulting opsin sequences from Sparkling Lake were used as references for mapping Sparkling metatranscriptome reads. For Mendota and Trout Bog, the resulting GEODES opsin sequences from the above steps were compared to opsin sequences longer than 600 bp from the Mendota and Trout Bog combined assembly respectively by BLASTN to identify GEODES sequences not represented in the corresponding combined assembly as defined by <97% nucleotide identity. The identified GEODES sequences were added to the >600 bp opsin sequences from the corresponding combined assembly to generate the mapping reference for Mendota and Trout Bog respectively. Metatranscriptome reads used for mapping were previously processed to remove adapter sequences, low-quality or short reads, and rRNA sequences by Linz *et al*. (2020), and were mapped to corresponding opsin references with the internal standard sequence by using bowtie2 (v2.2.2) with default settings. Metatranscriptomes with low read counts from the internal standard were discarded according to Linz *et al*. (2020). Opsin transcript read counts were normalized using the read count from the internal standard and converted to transcript concentrations. RAIN test (Thaben & Westermark 2014) was conducted to detect genes with diel cyclic trends with *p*-values of less than 0.05 for cyclic genes.

## RESULTS

### 1. Diversity of freshwater opsin genes

#### Overview

Here, we studied opsin gene diversity using three combined metagenome assemblies, generated by co-assembling reads from all metagenomes within the ME (Mendota Epilimnion, 94 metagenomes), TE (Trout Bog Epilimnion, 45 metagenomes), and TH (Trout Bog Hypolimnion, 45 metagenomes) groups (Linz *et al*. 2018). Altogether, from the three combined assemblies, opsin genes were recovered in 42 of 184 MAGs (**Table 1**) plus 50 bins that did not pass our criteria for quality MAGs. Genome completeness/contamination assessments and taxonomic classification of these MAGs and bins, and their opsin gene IMG IDs are provided in **Table S5**. At the phylum level, the classification of these opsin genes through classifying their resident MAG/bins was largely consistent with the IMG classification of opsin gene-containing contigs based on the consensus BLASTP hits for all genes on the contig (**Table S5**). Therefore, we assumed that opsin genes from these MAGs/bins were suitable phylogenetic anchors to classify metagenome opsin sequences in un-binned contigs to at least the phylum level.

**Table 1.**
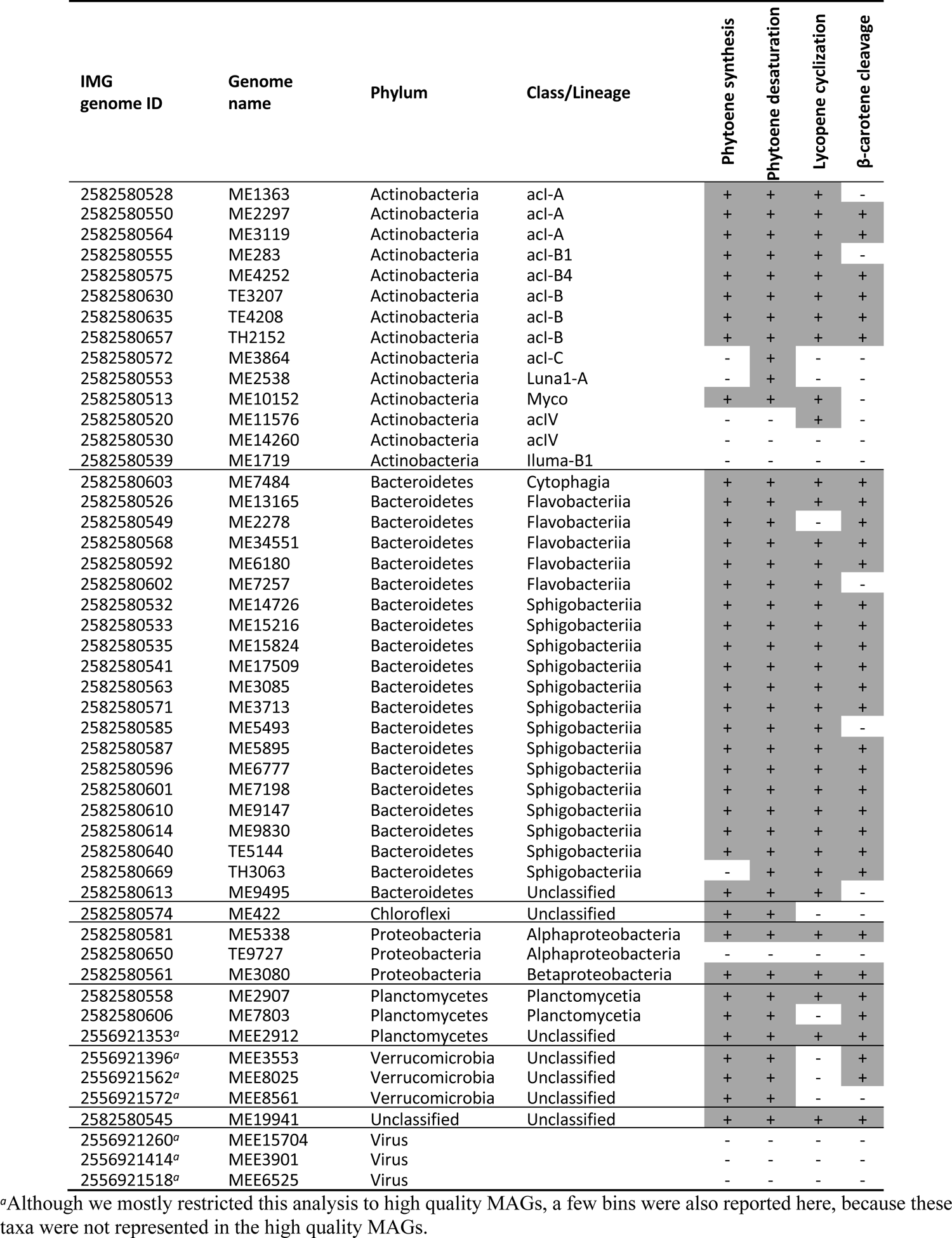
Mendota and Trout Bog MAGs containing opsin genes and the occurrence of key genes in their retinal biosynthesis pathway

| IMG genome ID | Genome name | Phylum | Class/Lineage | Phytoene synthesis | Phytoene desaturation | Lycopene cyclization | β-carotene cleavage |
| --- | --- | --- | --- | --- | --- | --- | --- |
| 2582580528 | ME1363 | Actinobacteria | acI-A | + | + | + | - |
| 2582580550 | ME2297 | Actinobacteria | acI-A | + | + | + | + |
| 2582580564 | ME3119 | Actinobacteria | acI-A | + | + | + | + |
| 2582580555 | ME283 | Actinobacteria | acI-B1 | + | + | + | - |
| 2582580575 | ME4252 | Actinobacteria | acI-B4 | + | + | + | + |
| 2582580630 | TE3207 | Actinobacteria | acI-B | + | + | + | + |
| 2582580635 | TE4208 | Actinobacteria | acI-B | + | + | + | + |
| 2582580657 | TH2152 | Actinobacteria | acI-B | + | + | + | + |
| 2582580572 | ME3864 | Actinobacteria | acI-C | - | + | - | - |
| 2582580553 | ME2538 | Actinobacteria | Luna1-A | - | + | - | - |
| 2582580513 | ME10152 | Actinobacteria | Myco | + | + | + | - |
| 2582580520 | ME11576 | Actinobacteria | acIV | - | - | + | - |
| 2582580530 | ME14260 | Actinobacteria | acIV | - | - | - | - |
| 2582580539 | ME1719 | Actinobacteria | Iluma-B1 | - | - | - | - |
| 2582580603 | ME7484 | Bacteroidetes | Cytophagia | + | + | + | + |
| 2582580526 | ME13165 | Bacteroidetes | Flavobacteriia | + | + | + | + |
| 2582580549 | ME2278 | Bacteroidetes | Flavobacteriia | + | + | - | + |
| 2582580568 | ME34551 | Bacteroidetes | Flavobacteriia | + | + | + | + |
| 2582580592 | ME6180 | Bacteroidetes | Flavobacteriia | + | + | + | + |
| 2582580602 | ME7257 | Bacteroidetes | Flavobacteriia | + | + | + | - |
| 2582580532 | ME14726 | Bacteroidetes | Sphigobacteriia | + | + | + | + |
| 2582580533 | ME15216 | Bacteroidetes | Sphigobacteriia | + | + | + | + |
| 2582580535 | ME15824 | Bacteroidetes | Sphigobacteriia | + | + | + | + |
| 2582580541 | ME17509 | Bacteroidetes | Sphigobacteriia | + | + | + | + |
| 2582580563 | ME3085 | Bacteroidetes | Sphigobacteriia | + | + | + | + |
| 2582580571 | ME3713 | Bacteroidetes | Sphigobacteriia | + | + | + | + |
| 2582580585 | ME5493 | Bacteroidetes | Sphigobacteriia | + | + | + | - |
| 2582580587 | ME5895 | Bacteroidetes | Sphigobacteriia | + | + | + | + |
| 2582580596 | ME6777 | Bacteroidetes | Sphigobacteriia | + | + | + | + |
| 2582580601 | ME7198 | Bacteroidetes | Sphigobacteriia | + | + | + | + |
| 2582580610 | ME9147 | Bacteroidetes | Sphigobacteriia | + | + | + | + |
| 2582580614 | ME9830 | Bacteroidetes | Sphigobacteriia | + | + | + | + |
| 2582580640 | TE5144 | Bacteroidetes | Sphigobacteriia | + | + | + | + |
| 2582580669 | TH3063 | Bacteroidetes | Sphigobacteriia | - | + | + | + |
| 2582580613 | ME9495 | Bacteroidetes | Unclassified | + | + | + | - |
| 2582580574 | ME422 | Chloroflexi | Unclassified | + | + | - | - |
| 2582580581 | ME5338 | Proteobacteria | Alphaproteobacteria | + | + | + | + |
| 2582580650 | TE9727 | Proteobacteria | Alphaproteobacteria | - | - | - | - |
| 2582580561 | ME3080 | Proteobacteria | Betaproteobacteria | + | + | + | + |
| 2582580558 | ME2907 | Planctomycetes | Planctomycetia | + | + | + | + |
| 2582580606 | ME7803 | Planctomycetes | Planctomycetia | + | + | - | + |
| 2556921353<sup>a</sup> | MEE2912 | Planctomycetes | Unclassified | + | + | + | + |
| 2556921396<sup>a</sup> | MEE3553 | Verrucomicrobia | Unclassified | + | + | - | + |
| 2556921562<sup>a</sup> | MEE8025 | Verrucomicrobia | Unclassified | + | + | - | + |
| 2556921572<sup>a</sup> | MEE8561 | Verrucomicrobia | Unclassified | + | + | - | - |
| 2582580545 | ME19941 | Unclassified | Unclassified | + | + | + | + |
| 2556921260<sup>a</sup> | MEE15704 | Virus |  | - | - | - | - |
| 2556921414<sup>a</sup> | MEE3901 | Virus |  | - | - | - | - |
| 2556921518<sup>a</sup> | MEE6525 | Virus |  | - | - | - | - |

<sup>a</sup>Although we mostly restricted this analysis to high quality MAGs, a few bins were also reported here, because these taxa were not represented in the high quality MAGs.

We constructed a phylogenetic tree with previously published references and opsin sequences longer than 200 amino acids in the three combined assemblies, which include sequences from MAGs/bins and un-binned contigs (**Figure 1**, with a rectangular view of this tree with more detailed gene information in **Figure S1**). Metagenome opsin sequences shorter than 200 amino acids were aligned and added to the phylogenetic tree (**Figure S2**). Based on the coverage-weighted opsin gene abundance in the assemblies, we estimated the percent distribution of opsin genes among different taxa and rhodopsin types in each lake/layer (**Figure S3**).

**Figure 1.**
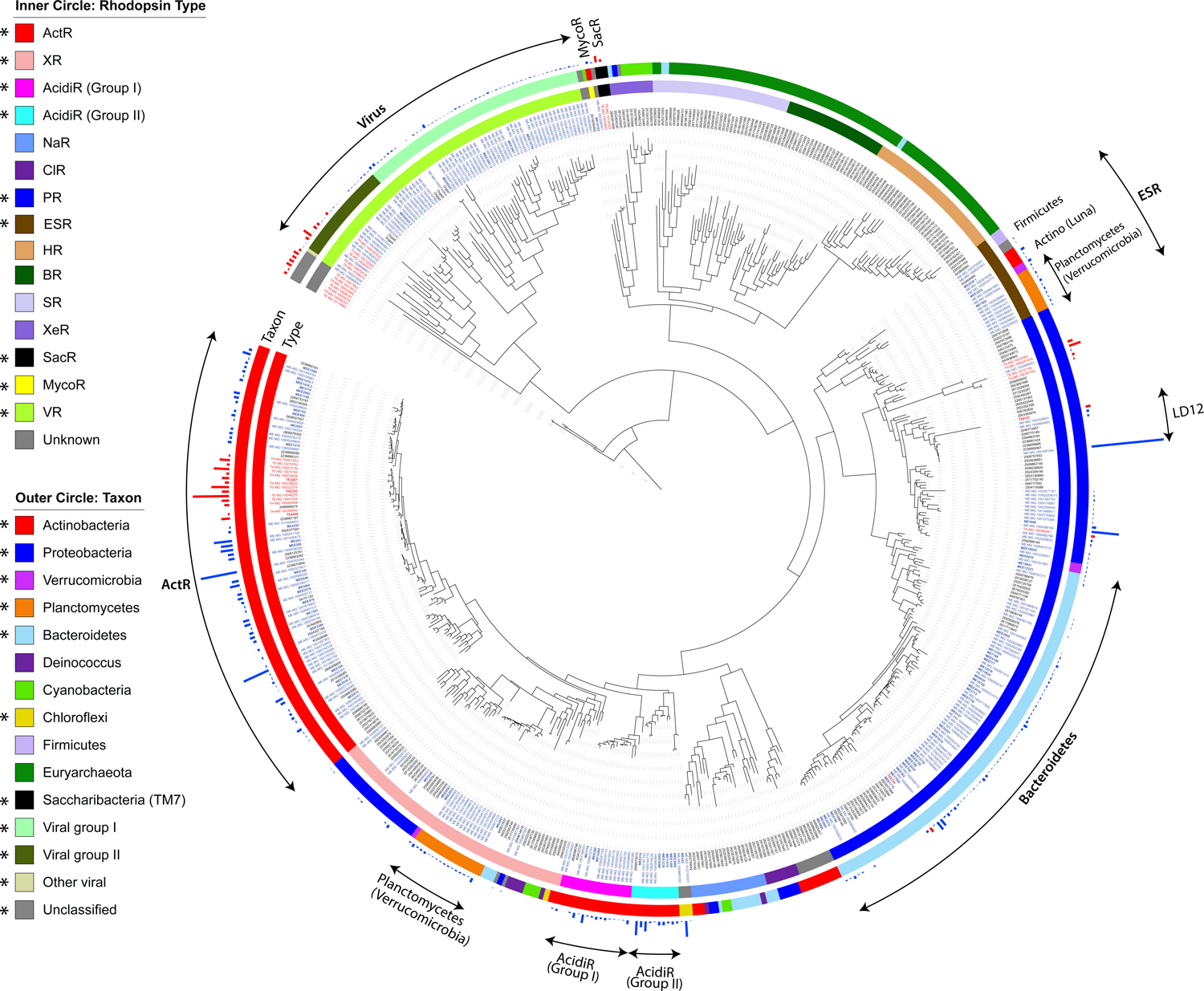
Phylogenetic tree of opsin protein sequences. This tree was viewed and annotated using the Interactive Tree of Life tool (http://itol.embl.de). Opsin gene names are colored as following: blue for ME, red for TE and TH metagenomes, and black for references from the database. For sequences in MAGs/bins (highlighted with bold fonts), gene names start with “ME”, “MEE”, “TE”, and “TH” respectively. For the rest (un-binned) metagenome sequences, gene names start with “ME-MG”, “TE-MG”, and “TH-MG”, respectively. The inner circle color strip indicates Rhodopsin types, and the outer circle color strip indicates taxon. Rhodopsin types and taxon present in our metagenomes were indicated with “*” in the legend. The outmost bar indicates normalized coverage-weighted abundance of opsin genes in the metagenome (blue for ME, and red for TE and TH). Some branches were labeled to highlight points discussed in the manuscript. Rhodopsin types include ActR (actinorhodopsin), XR (xanthorhodopsin), AcidiR (acidirhodopsin), NaR (sodium-pumping rhodopsins), CIR (chloride-pumping rhodopsins), PR (proteorhodopsin), ESR (*Exiguobacterium sibiricum* rhodopsin), HR (halorhodopsin), BR (bacteriorhodopsin), SR (sensory rhodopsin), XeR (xenorhodopsin), SacR (Saccharibacteria rhodopsin), MycoR (DTG rhodopsins represented by the freshwater Myco tribe), and VR (viral rhodopsin).

The majority of recovered opsin sequences belong to known H^+^-pumping branches, such as actinorhodopsin (ActR), proteorhodopsin (PR), xanthorhodopsin (XR) and *Exiguobacterium sibiricum* rhodopsin (ESR) (**Figure 1**). Most sequences have the conserved DTE (aspartic acid, threonine, glutamic acid) motif corresponding to Positions 85, 89, and 96 (BR numbering) for H^+^ accepting and donation (Beja & Lanyi 2014). The spectral tuning residue on the recovered opsin sequences is a hydrophobic residue methionine (M) or leucine (L) for absorbing green light, with a few exceptions that have the hydrophilic glutamine (Q) for absorbing blue light (Gómez-Consarnau *et al*. 2007; Man *et al*. 2003). Therefore the majority of recovered opsin genes were predicted to encode green light-absorbing H^+^ pumps. In Mendota, opsin genes were recovered from all major freshwater phyla except Cyanobacteria. These included Actinobacteria, Bacteroidetes, Proteobacteria, Planctomycetes, Verrucomicrobia, and Chloroflexi, and the first three phyla were the major contributors to opsin gene abundance (**Figure S3**). Whereas in Trout Bog, opsin genes were restricted to fewer taxa, mainly from Actinobacteria (acI-B2 in particular) and some from Proteobacteria and *Candidatus* Saccharibacteria (TM7) (**Figure S3**).

As we only recovered a small number of opsin sequences in Trout Bog MAGs (**Table S6**), we also searched the 12665 high-quality MAGs from the stratfreshDB which was generated from 41 freshwater lakes and ponds, including many “clearwater” (not stained brown) and brown-water humic environments. We found a total of 2011 opsin genes, most of which were also predicted to encode green light-absorbing H^+^-pumps based on the DTE motif (**Figure 2a**). Similarly, Actinobacteria, Bacteroidetes and Proteobacteria were the major opsin-containing phyla (**Table S7**). In addition to the phyla found in Mendota and Trout Bog, a very small number of opsin genes were also recovered from phyla such as Acidobacteriota and Bdellovibrionota from clearwater environments. Like the contrast between Mendota and Trout Bog (**Figure 3a**), opsin genes occurred more frequently in clearwater MAGs than in humic water MAGs from the stratfreshDB (**Table S7**).

**Figure 2.**
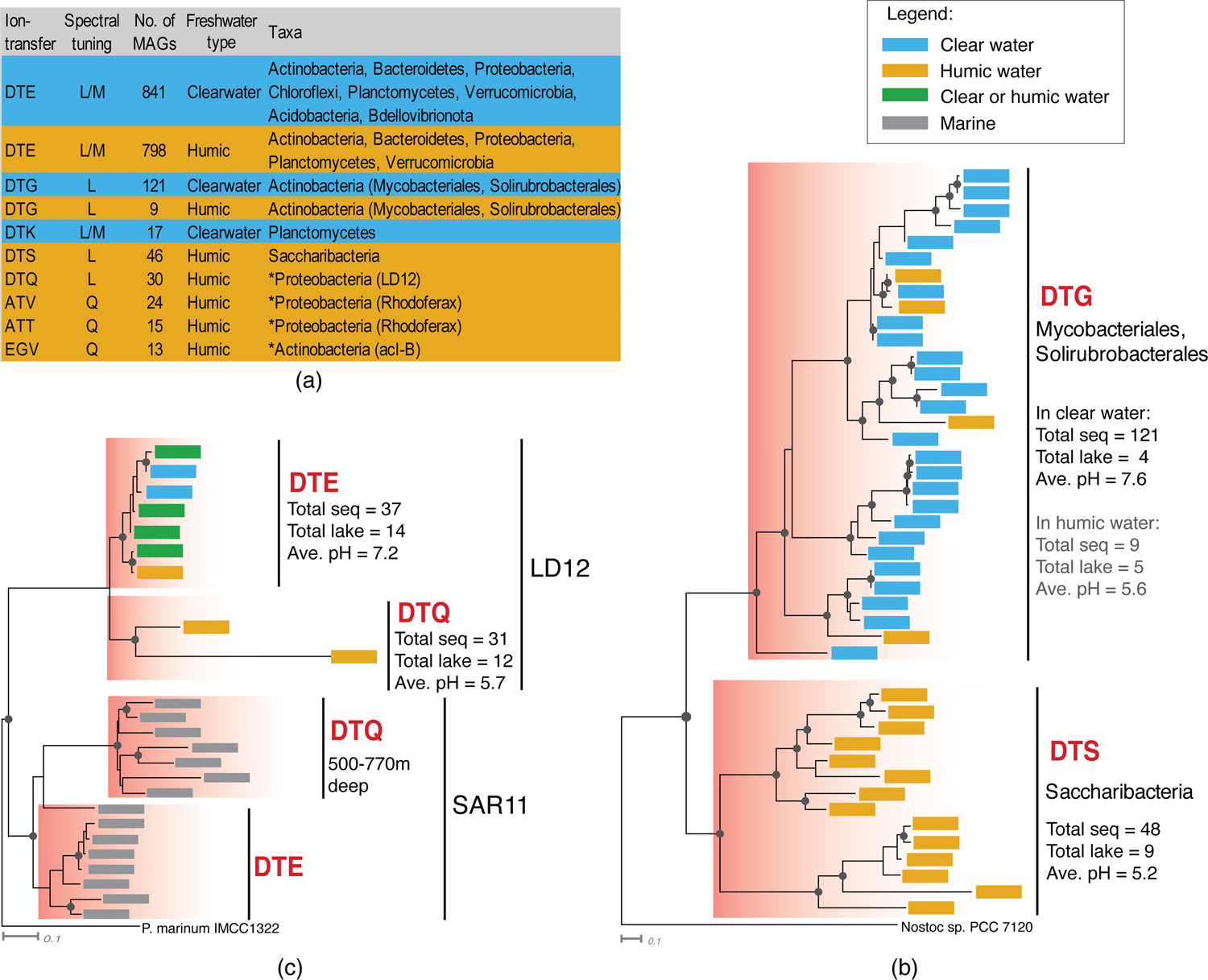
Variants of the cytoplasmic ion-transferring motif. (**a**) Distribution of major variants of the ion-transferring motif, listing together with the spectral-tuning residue, the number of opsin sequences with that motif from Mendota, Trout Bog and the stratfreshDB MAGs, the occurrence of these MAGs in clear or humic waters, and the taxa that contributed to these variants. The asterisk symbol ahead of the taxa name indicates that the DTE motif is also present in members of that taxa (from different MAGs). (**b**) Phylogenetic tree of opsins from LD12 and SAR11, with *Ca.* Puniceispirillum marinum (SAR116) as the outgroup. (**c**) Phylogenetic tree of opsins from the MycoR and SacR groups (DTG/DTS cluster), with a xenorhodopsin from *Nostoc* sp. as the outgroup. In both (**b**) and (**c**), sequences from Mendota, Trout Bog and the stratfreshDB MAGs were clustered based on 99% amino acid identity, and representative sequences were used to construct maximum likelihood phylogenetic trees using PhyML 3.0, with the LG substitution model and the gamma distribution parameter estimated by PhyML. Nodes labeled with a solid circle have bootstrap support values higher than 50%. For freshwater clades, the total number of sequences, the total number of their occurring lakes and the average pH of their occurring environments were listed for each motif variant-defined group. For the DTG group, as most (121 out of 130) sequences were from four clearwater lakes, these values listed for clear and humic waters respectively.

**Figure 3.**
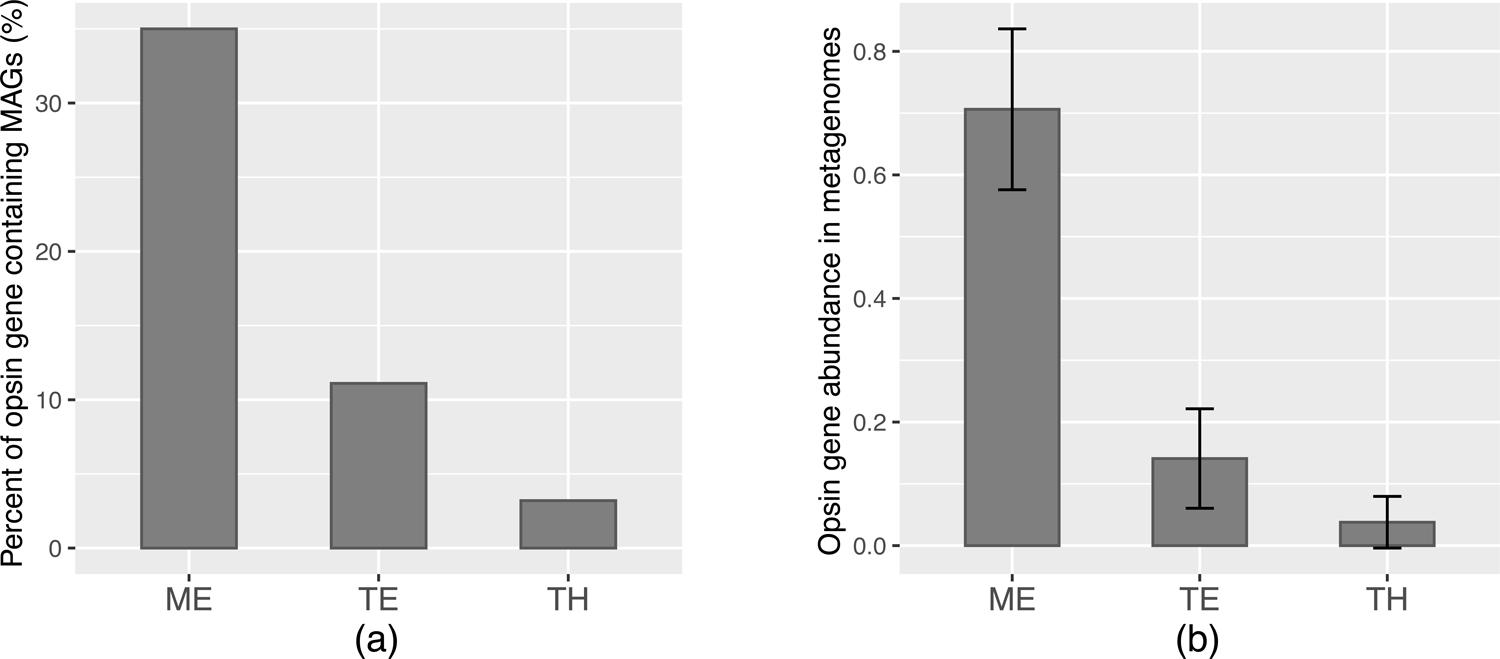
The significance of opsin genes in ME, TE and TH. (**a**) Percentage of opsin gene containing MAGs among the total of 103, 36, and 63 bacterial MAGs recovered in ME, TE and TH metagenomes, respectively. (**b**) Average of opsin gene relative abundance in metagenomes estimated by opsin gene coverage normalized by the coverage of single-copy genes. Error bars indicate standard deviations derived from the 95, 29 and 29 individual metagenomes generated from ME, TE and TH (listed in Table S3).

### Opsin gene prevalence and diversity in freshwater Actinobacteria

Actinobacteria was the phylum possessing the largest number of opsin genes, accounting for 46-61% of all opsin gene coverage in each of the three combined assemblies (**Figure S3**). In ME, opsin genes were widely distributed in nearly all lineages/clades of Actinobacteria that were previously known to harbor opsin genes (**Figure S4**). The acI lineage, particularly Ca. Planktophila (acI-A) and Nanopelagicus (acI-B), was the most prominent contributor to opsin genes in ME (**Figure S3**). Although acI was also the most abundant contributor to opsin genes in TE and TH, their ActR genes were confined within a branch of acI-B, likely from acI-B2. This branch exclusively contained opsin sequences from humic lakes, consistent with the finding that acI-B2 prefers humic lakes as compared to acI-B1 (Garcia *et al*. 2014; Newton *et al*. 2006).

Most Actinobacterial opsin genes belong to ActR, an outward H^+^-pumping group phylogenetically close to XR. Subgroup I of XR also has a secondary chromophore, a noncovalently bound carotenoid functioning as a light-harvesting antenna, in addition to the retinal (Balashov *et al*. 2005; Béjà *et al*. 2000; Imasheva *et al*. 2009; Vollmers *et al*. 2013). Notably, nearly all ActR sequences recovered in our study contain the signature glycine that replaces a bulky residue on helix 5, which permits binding of the carotenoid antenna salinxanthin in the well-characterized XR from *Salinibacter ruber* (Balashov *et al*. 2005). Whether any individual ActR-encoding microbe synthesizes a carotenoid antenna and whether particular ActR exemplars bind a secondary carotenoid may need to be addressed on a case-by-case basis but the presence of the signature glycine residue and the genetic machinery for a secondary carotenoid biosynthesis have been noted (Dwulit-Smith *et al*. 2018), and in at least one case the binding of a native actinobacterial carotenoid to an ActR has been demonstrated (Chuon *et al*., 2021).

In addition to ActR, a rhodopsin group phylogenetically distinct from the typical ActR was previously discovered in both freshwater and marine actinobacterial MAGs belonging to the order of Acidimicrobiales and was named acidirhodopsin (short as “AcidiR” in our study in contrast to ActR) (Ghai *et al*. 2014; Mizuno *et al*. 2015). According to the freshwater taxonomy, the AcidiR-containing freshwater MAG (Ghai *et al*. 2014) belongs to the Iluma-B clade (alternative name acIV-B). Here in ME, we also found sequences belonging to this Iluma-B AcidiR clade (**Figures 1** and **S4**), accounting for ∼5% of all opsin gene coverage in the combined ME assembly (**Figure S3**). This AcidiR clade did not form a monophyletic group with ActR, but instead was a sister clade to the branch formed by ActR and XR (**Figures 1** and **S1**), similar to its placement in previously reconstructed opsin phylogenetic trees (Cabello-Yeves *et al*. 2017; Durán-Viseras *et al*. 2019; Mizuno *et al*. 2015). In addition, we identified a different branch of opsin sequences belonging to Iluma (likely Iluma-A), accounting for ∼6% of all opsin gene coverage in the combined ME assembly. This branch did not form a monophyletic clade with the previously identified AcidiR branch (**Figures 1 and S1**). Therefore, we use “AcidiR (Group I)” and “AcidiR (Group II)” to distinguish them. A difference between these two groups is that the conserved proline in the middle of helix 4 of ActR is also conserved in AcidiR Group I but missing in Group II (as well as XR). Such a proline residue may impact opsin structure and thus render different photoproperties as previously postulated (Dwulit-Smith *et al*. 2018). Notably, the bootstrap values associated with the relative placement among the ActR, XR and AcidiR branches were lower than 50%, and this was also the case in other studies (Durán-Viseras *et al*. 2019; Mizuno *et al*. 2015). As such, the phylogeny and evolutional history among these clades needs to be further resolved.

**Figure 4.**
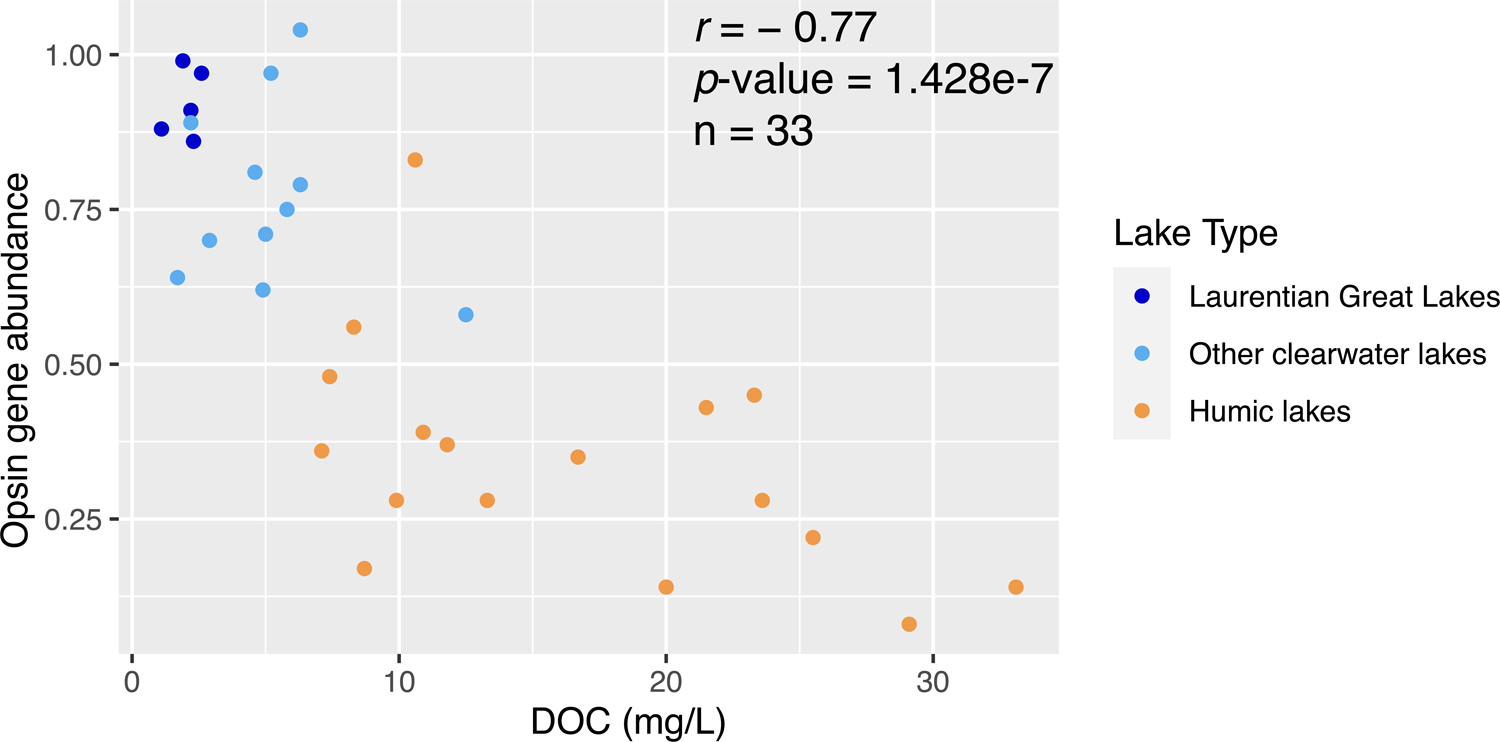
Opsin gene relative abundance in the surface layer of the 33 lakes versus the lake DOC levels. The 33 lakes, DOC levels, Accession No. of metagenomes included in this analysis and corresponding references for published studies were listed in Table S2. For lakes with multiple metagenomes (derived from different dates, depths, or studies), opsin gene abundance was estimated from individual metagenomes from the surface layer and averaged for that lake. Among the 33 lakes, four small boreal lakes from Buck *et al*. (2021) only had total organic carbon (TOC) measurements. As the difference between DOC and TOC was generally small for lakes in that study, we use TOC in place of DOC for those four lakes on this plot.

Besides ActR and AcidiR, an ME MAG classified to the Myco tribe within the acTH2 lineage of Actinobacteria contains an opsin gene that has an unusual H^+^-pumping motif, DTG (G for glycine), forming a clade with another ME sequence with the DTG motif. This clade is phylogenetically closer to the sensory inward H^+^-pumping xenorhodopsins (XeR) than to other well-studied rhodopsin groups (**Figures 1** and **S1**). We tentatively named it “MycoR” as it was identified in the Myco tribe. Many more DTG opsin sequences belonging to this cluster were found in numerous stratfreshDB MAGs classified to Mycobacteriales and Solirubrobacterales within Actinobacteria, and notably most of these MAGs were recovered from clearwater environments (**Figures 2a** and **2b**). DTG rhodopsins were initially discovered in soil- and plant-associated Proteobacteria (Harris *et al*. 2015; Sudo & Yoshizawa 2016; Suzuki *et al*. 2022) and our findings expand the taxonomic diversity of DTG rhodopsins.

### Wider than previously thought opsin gene distribution in Bacteroidetes

Actinobacteria have long been regarded as the major phylum containing opsin genes among freshwater bacteria. For example, among single cells recovered from freshwater lakes, the six opsin-containing Sphingobacteriia SAGs were dwarfed by the large number (∼100) of opsin-containing Actinobacteria SAGs (Martinez-Garcia *et al*. 2012). In our study, opsin genes from Bacteroidetes were exclusively within the PR group, and correspond to MAGs placed in the Sphingobacteriia, Flavobacteriia, and Cytophagia classes (**Table 1** and **Figure S1**). They were mostly from low-abundance populations in ME (**Figure 1**), yet when added together, they accounted for ∼13% of all opsin gene coverage in the combined ME assembly (**Figure S3**). Notably, in both Mendota and clear waters from the stratfreshDB Actinobacteria and Bacteroidetes had the highest occurrences of opsin-containing MAGs among all MAGs (**Tables S6** and **S7**), suggesting that, similar to Actinobacteria, Bacteroidetes in clear waters also widely possess opsin genes, and therefore, could be a previously overlooked photoheterotrophic lineage in clearwater lakes.

### Opsin gene diversity in Proteobacteria

Proteobacteria are abundant and prevalent in freshwater, which is also reflected by the large numbers of Proteobacteria MAGs recovered from Mendota, Trout Bog and the stratfreshDB dataset. Yet compared to Actinobacteria and Bacteroidetes, relatively fewer opsin sequences were recovered from this phylum, especially in humic water (**Tables S6** and **S7**). For example, the ubiquitous *Polynucleobacter* spp., which frequently occur in humic water, mostly lack opsin genes. Freshwater proteobacterial opsin genes in our study belong to the PR and XR groups (**Figures 1** and **S3**). Notably, several un-binned metagenome sequences in the PR branch, including one occurring at an extremely high abundance, fall within the freshwater *Candidatus* Fonsibacter (LD12) (**Figures 1** and **S1**). LD12 is a sister clade of marine SAR11, which is the major contributor to rhodopsins in the ocean (Maresca *et al*. 2018; Olson *et al*. 2018; Sieradzki *et al*. 2018). As LD12 are very abundant in Mendota, opsin genes in the LD12 clade alone contributed to nearly 11% of all opsin gene coverage in the combined ME assembly. Among the stratfreshDB MAGs, besides LD12, ubiquitous freshwater genera such as *Limnohabitans*, *Rhodoferax*, and “*Ca*. Methylopumilus” were major bacterial taxa with opsin genes. Most proteobacterial opsins have the DTE motif for pumping H^+^, especially in clearwater MAGs. Members of a few taxa such as LD12 and *Rhodoferax* have ion-pumping motifs other than DTE in humic water (**Figure 2a**), although the typical DTE motif also occurred in other members of these taxa. For example, some humic water LD12 MAGs have the DTQ (Q for glutamine) motif (**Figure 2c**). Proteobacterial opsins with such unusual motifs were almost exclusively from humic water, accounting for nearly 40% of all proteobacterial opsin sequences from humic water MAGs in the stratfreshDB, and therefore their importance in humic aquatic environments is unlikely to be negligible.

### Opsin genes in other bacterial taxa

Opsin genes belonging to Planctomycetes, Verrucomicrobia, and Chloroflexi were distributed in the PR, XR and ESR branches (**Figure 1**). ESR, a branch phylogenetically closest to PR, was named after the firmicute *Exiguobacterium sibiricum* which has an unusual H^+^-translocating motif, DTK (K for lysine) (Balashov *et al*. 2013). The ESR branch includes Firmicutes and freshwater Planctomycetes, Verrucomicrobia, and Actinobacteria Luna1 and Luna2 (**Figures 1** and **S1**). Like in *Exiguobacterium* spp., Planctomycetes sequences in the ESR branch also have the DTK motif, and notably these planctomycetes ESR sequences were recovered from clearwater environments (**Figure 2a**), although planctomycetes with the DTE motif in other branches were recovered from both clear and humic waters.

Opsin genes belonging to *Candidatus* Saccharibacteria (previously candidate division TM7) were recovered from un-binned contigs in Trout Bog. Their sequences are highly similar (∼93% amino acid identity) to the opsin gene in a genome bin classified as TM7 (TM7 FNED7-bin-18, IMG genome ID 2648501560) recovered in an enrichment culture originating from the bog-like basin of Grosse Fuchskuhle in Germany (**Figure S1**) (Garcia *et al*. 2014). In fact, Saccharibacteria rhodopsins (SacR) were also recently discovered from stratfreshDB MAGs and function as a light-driven outward H^+^ pump (Jaffe *et al*. 2022). We noted that all the 46 SacR sequences from the stratfreshDB also exclusively occurred in humic waters. SacR have the DTS (S for serine) H^+^-pumping motif and with MycoR, they form a DTG/DTS cluster (**Figures 1**, **2b, S1**).

### Viral opsin genes

Viral opsin genes were first discovered in giant viruses that infect marine unicellular eukaryotes, such as Phycodnaviruses and *Phaeocystis globosa* virus in Organic Lake, an Antarctic lake with extremely salty water (Yutin & Koonin 2012). They were later detected in the Red Sea and were referred to as “VR” for viral rhodopsins (Philosof & Beja 2013). Protein structure characterization and functional analyses revealed that they encode light-driven H^+^-pumps (Needham *et al*. 2019) or light-gated ion channels (Bratanov *et al*. 2019; Zabelskii *et al*. 2020). In our study, we recovered a large opsin gene cluster outside of all bacterial and archaeal branches in the phylogenetic tree (**Figures 1** and **S1**). Three viral bins (see **Supplementary Text** and **Tables S8** and **S9** for their classification as viruses) recovered from ME each contain one to two copies of opsin genes belonging to this cluster (**Tables 1** and **S5**). This cluster also harbored opsin sequences from algal viruses previously recovered from marine or salty water (**Figure S1**), supporting the viral origin of this cluster. These freshwater viral opsin genes also form two distinct sub-clusters, Viral Group I and II as previously defined (Yutin & Koonin 2012), each harboring both freshwater and marine sequences (**Figure S1**). The function of these freshwater viral rhodopsins, their roles in virus-host interactions, and their influences on host physiology and ecology are yet to be studied.

### 2. Retinal biosynthesis

Retinal is a photosensitive chromophore cofactor of rhodopsin. One important question is whether the opsin gene-containing genomes we recovered also have the genetic potential for retinal biosynthesis to form functional rhodopsin. For this, we checked key genes responsible for the last few steps in retinal biosynthesis, including phytoene synthesis, phytoene desaturation, lycopene cyclization, and β-carotene cleavage. We found many opsin-containing MAGs also have the genetic potential for retinal biosynthesis (**Table 1**). Due to incompleteness, it is difficult to conclude the lack of genes based on MAGs. However, some patterns are consistent. For example, all viral bins lacked retinal biosynthesis genes, and therefore, they might rely on their hosts or the environment for the chromophore. In addition, among Actinobacteria, retinal biosynthesis genes were mostly found within acI, particularly acI-A and acI-B. This trend is supported by a total of 74 freshwater Actinobacteria genomes, including SAGs and MAGs from other studies that span most freshwater Actinobacteria lineages (**Table S10**). Although opsin genes are widely distributed among these lineages (except acV), oxygenase genes responsible for β-carotene cleavage are only present in some tribes within the acI-A and acI-B clades (**Table S10**). Thus, Actinobacteria without the retinal biosynthesis capability need to take up retinal from the environment if their opsins are to be photoactive. For example, the freshwater Actinobacterium *Rhodoluna lacicola* WH-Ta8^T^ conducted light-induced H^+^ pumping only after exogenous retinal was provided (Keffer *et al*. 2015).

### 3. Ecological distribution of opsin genes

We estimated the relative abundance of opsin genes in metagenomes by comparing the coverage-weighted abundance of opsin genes to single-copy essential housekeeping genes that have comparable lengths to opsin genes. As opsin genes in our MAGs are mostly single-copy (**Table S5**) and viral contribution to the total opsin genes is relatively small (**Figure S3**), such estimation largely reflects the fraction of bacteria that possess opsin genes in the total community. We performed this estimation using individual metagenomes generated from each time point from ME, TE, and TH. Comparison between the two epilimnia indicated that opsin gene abundance was higher in ME (∼70%) than in TE (∼14%) (**Figure 3b**). We therefore hypothesized that opsin gene distribution could be influenced by lake DOC and optical properties, in particular, that high DOC and light-absorbing humic substances and thus low light availabilities can lead to the low occurrence of opsin genes, whereas low DOC and good light penetration may enrich for opsin genes.

### Comparison among the surface layers of 33 lakes

To test this hypothesis, we included a total of 33 freshwater systems for which metagenome data and DOC levels are publicly available (**Table S2**). These lakes range from oligotrophic, mesotrophic, and eutrophic clearwater (not stained brown) lakes, to dystrophic brown-water humic lakes, spanning a wide range of DOC levels and exhibiting different clarities. Examples of lake DOC levels, clarity as reflected in the Secchi depth and light diffuse attenuation coefficient, and other physicochemical and limnological characteristics are listed in **Table S1**. As opsin gene abundance can be influenced by water depth as discussed later, only metagenomes collected from the surface layer were included in this analysis to reduce variables. For lakes from the stratfreshDB, which largely consists of small boreal lakes, metagenomes collected within the first meter were included. An inverse correlation (Pearson correlation coefficient r = −0.77, *p-*value = 1.4e-7) between opsin gene abundance and DOC was observed (**Figure 4**). Opsin gene abundance was >50% in clearwater lakes irrespective of their trophic status. Especially, the abundance was extremely high (∼90%) in the Laurentian Great Lakes (Podowski *et al*. 2021) and Lake Lanier (Dam *et al*. 2020) where DOC levels are low (∼2 mg/L). Opsin genes were much less prominent in humic lakes with high terrestrial DOC. Extremely low abundance (<10%) was observed in the surface water of Lake Mekkojärvi, a highly humic boreal lake with ∼30 mg/L of DOC (Buck *et al*. 2021).

### Comparison at different depths within lakes

Within Trout Bog, comparing individual metagenomes sampled on the same day indicates the epilimnion had a higher opsin gene abundance than the hypolimnion, and the difference between layers is statistically significant (paired student T-test *p*-value < 1e-8, n = 29) (**Figure 3b**). Indeed, analyses of depth-discrete metagenome profiles show a steep decline of opsin gene abundance within the first few meters in humic lakes, leading to extremely low opsin gene levels in the deeper layer (**Figure 5**). By contrast, such a sharp decrease was not observed in clearwater lakes, where a depth-associated change in opsin gene abundance was much more moderate and slower (**Figure 5**).

**Figure 5.**
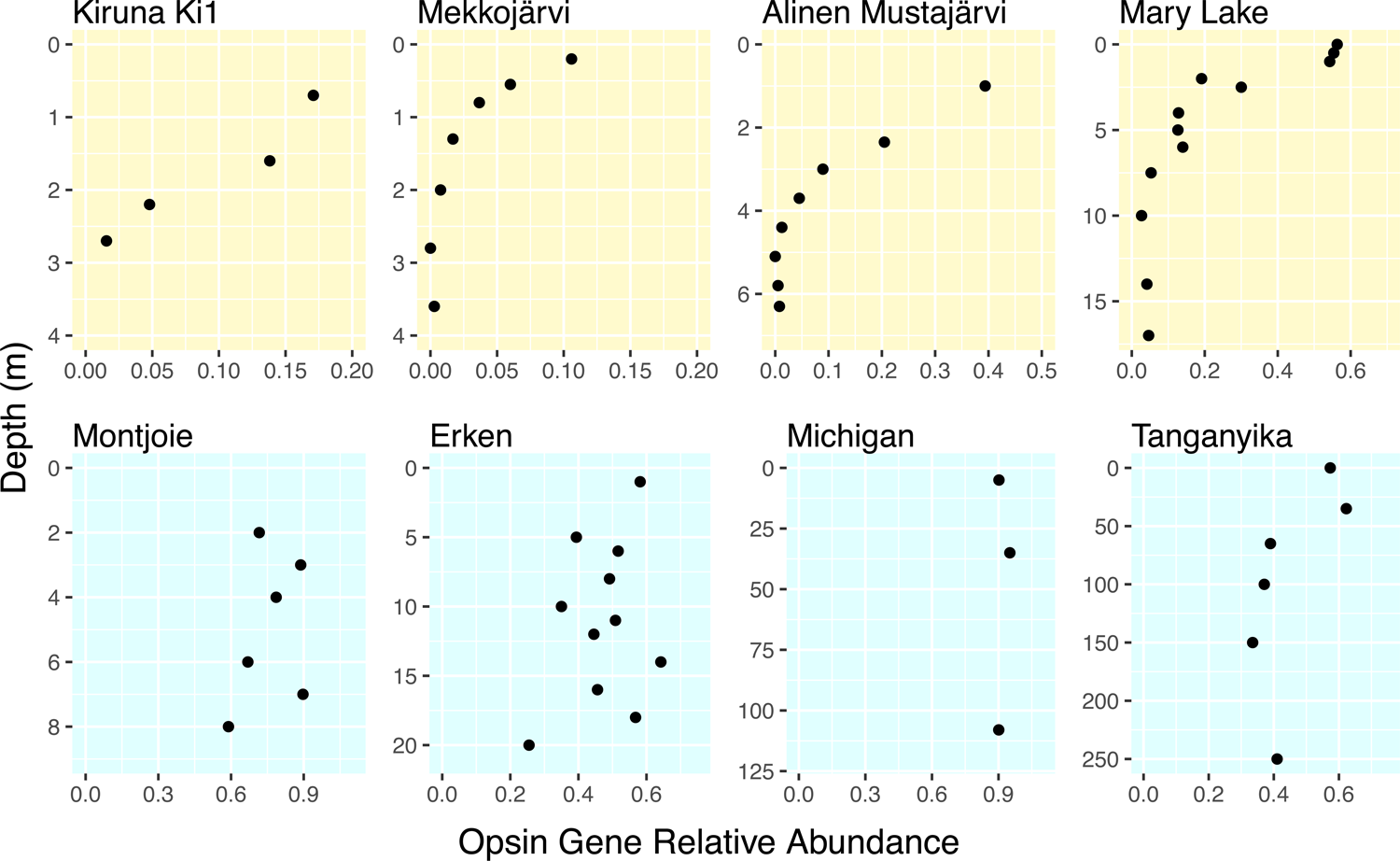
Depth profiles of opsin gene relative abundance. Lakes on the top panels are clearwater lakes, and on the bottom panels are humic lakes. For Lake Tanganyika, we only plotted abundances from the top 250 meters. Opsin gene abundance for samples at the depths of 300 m and 1200 m was 0.28 and 0.30 respectively. Metagenomes included in this analysis and corresponding reference papers were listed in Table S4.

### 4. Opsin gene expression and diel trends

We used metatranscriptomes previously generated from the epilimnia of Mendota, Trout Bog, and Sparkling Lake by Linz *et al*. (2020) to study opsin gene expression and diel cyclic rhythms. From metatranscriptome mapping outputs in the original analyses by Linz *et al*. (2020), the total opsin transcripts are about 0.5%, 0.3%, and 0.02% of the total mRNA for Mendota, Sparkling Lake, and Trout Bog respectively when averaged from all time points, and notably, a few opsin genes were among the top 500 most highly expressed genes within the entire metatranscriptomes in Mendota and Sparkling Lake. This indicates that opsin genes were highly expressed in these two clearwater lakes.

In our current analysis with more tailored and curated references for opsin genes, we found that 201 out of the 396 opsin genes in the Mendota opsin reference set had expression levels higher than the cutoff, and these numbers were 105 out of 173 and 18 out of 69 for Sparkling Lake and Trout Bog respectively. The total opsin transcripts per liter of lake water averaged from all time points showed that the total expression in the two clearwater lakes was about an order of magnitude higher than that in the humic bog (**Figure 6**). Bacterial phyla contributing to the most opsin expression were Actinobacteria, followed by Bacteroidetes and Proteobacteria in the two clearwater lakes, whereas opsin expression from Proteobacteria was minimal in the humic bog.

**Figure 6.**
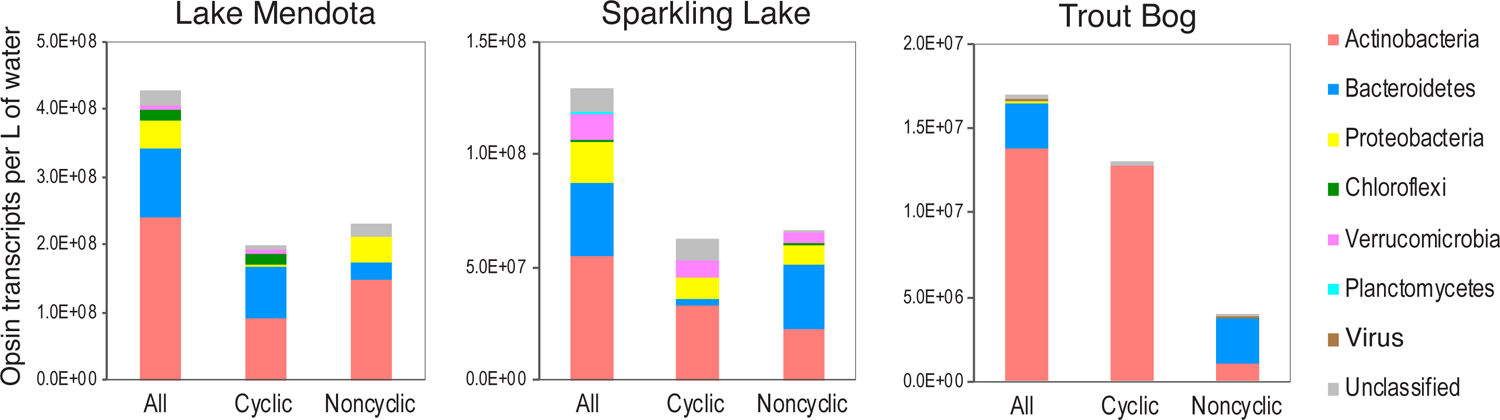
Total opsin transcripts per liter of lake water averaged from all time points included in this analysis. The values from Lake Mendota and Sparkling Lake are about an order of magnitude higher than that in Trout Bog.

Ranking individual opsin genes by their average transcript abundances indicated that opsin from Actinobacteria (mostly acI-A and acI-B), Bacteroidetes, and Proteobacteria (LD12 in particular) are among the most highly expressed opsins in the two clearwater lakes, whereas the most highly expressed opsins in Trout Bog were mainly from Actinobacteria acI-B (**Table S11**). Notably, both groups of AcidiR were expressed in the two clearwater lakes.

Among the expressed opsin genes, some exhibited diel trends. Depending on the photosynthetically active radiation (PAR) measured just below the surface of lake water, samples were categorized into “day” and “night” groups (Linz *et al*. 2020). Some taxa-specific diel patterns emerged. Cyclic genes from Actinobacteria tend to have peak expression at night, and Bacteroidetes opsins tend to peak at daytime in both Mendota and Sparkling Lake (**Figure 7**). In Trout Bog, only eight opsin genes were identified as cyclically transcribed. Seven of them belong to Actinobacteria and showed increased expression during the night and peaked before sunrise (**Figure S5**). However, Trout Bog results need further verification as only one day of data is available for this lake (Linz *et al*. 2020).

**Figure 7.**
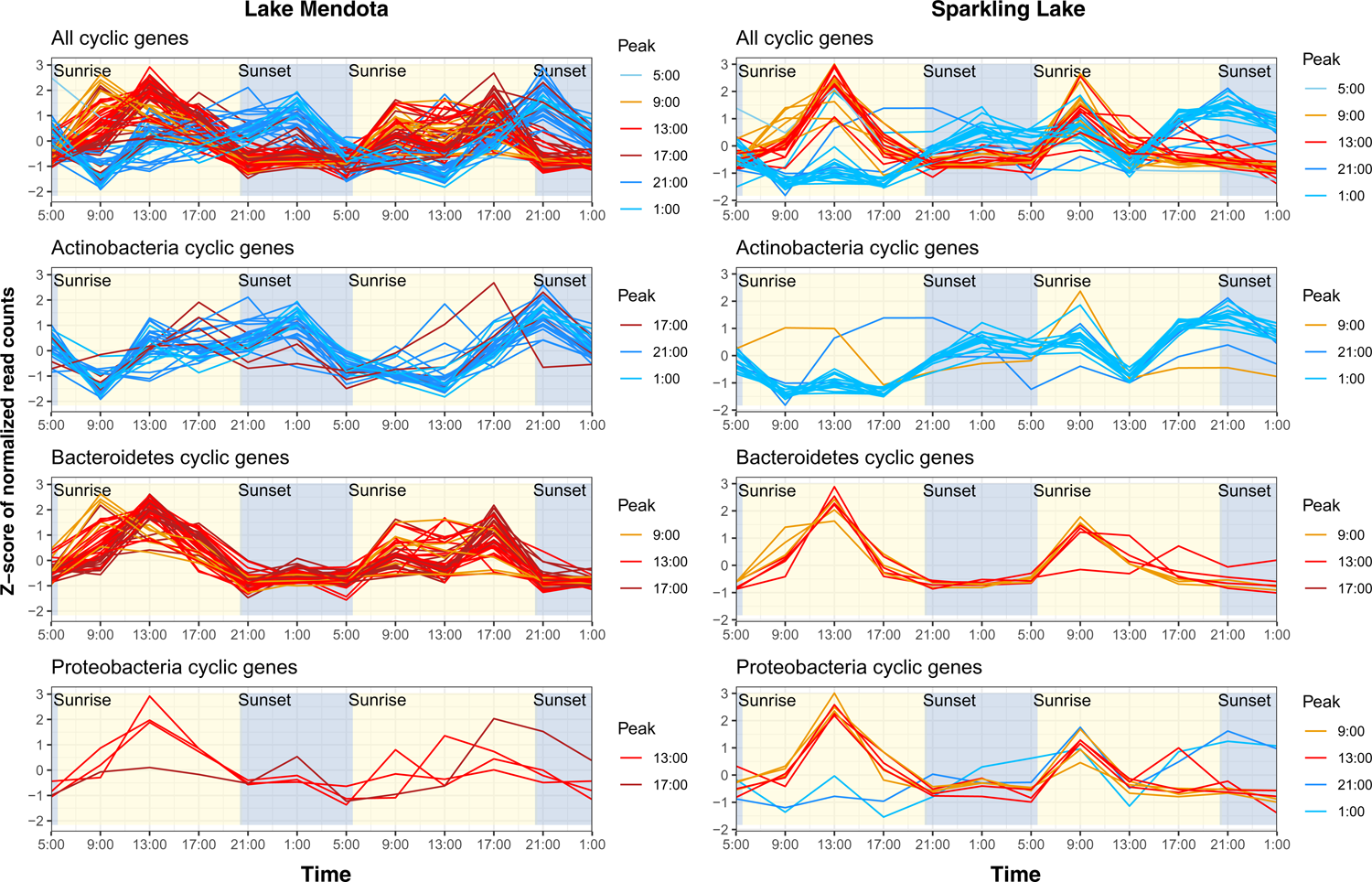
Diel cyclic trends of opsin gene expression. Cyclic genes were determined based on expression data collected from two consecutive days using the RAIN test with the *p*-value of < 0.05. For both lakes, the top panel shows all cyclic genes, and the bottom three panels show cyclic genes in Actinobacteria, Bacteroidetes and Proteobacteria respectively. Read counts in transcripts per liter were z-score transformed and are shown on y-axis. Genes were colored by the peak hour of their expression as determined in the RAIN test, with warm and cold colors for peaks at “day” and “night” (as determined by photosynthetically active radiation), respectively.

## DISCUSSION

### Diversity of opsin genes in clear and humic waters

Despite numerous studies on freshwater opsin genes, their diversity in brown-color, often acidic humic lakes had been largely unexplored, and the question remained about the difference, if any, between clear and humic waters with contrasting pH, DOC quantity/quality, and optical conditions.

Here we focused on the two main lakes which we have rich time-series metagenome data (and metatranscriptome data) to investigate the diversity and quantify the contribution of different taxa to the total opsin genes. Owing to the large number of MAGs from the stratfreshDB, we gained a more complete view of opsin diversity in both clear and humic waters. In general, findings from the stratfreshDB are consistent with what we observed with the Mendota and Trout Bog dataset. In both datasets, most opsin genes were predicted to encode green light-absorbing H^+^ pumps with the typical DTE variant for H^+^ donating and accepting, and Actinobacteria is the most abundant contributor of opsin genes in both humic and clearwater environments. Among the known taxa with opsin genes, we found wider than previously thought distribution within freshwater Bacteroidetes and identified novel clades, such as LD12 with the DTQ motif and MycoR. Our analysis also expanded the taxonomic spectra of freshwater opsins, especially in branches with unusual H^+^-pumping motifs.

Some differences between clear and humic waters emerged. Opsin genes were found more frequently in taxa from clear waters than in taxa from humic waters, which could lead to an observed low taxonomic diversity of opsin genes in a given humic lake, as we observed from Trout Bog. In addition, some opsins have non-typical H^+^-pumping motifs that might be associated with the two different lake types as discussed below (**Figure 2a**).

The DTG rhodopsin was previously recovered in soil- and plant-associated Proteobacteria as an light-driven outward H^+^-pump; and its photochemical behavior is pH dependent and H^+^-pumping slows down at high extracellular pH, which might prevent intracellular alkalization in alkalized environments (Harris *et al*. 2015; Sudo & Yoshizawa 2016; Suzuki *et al*. 2022). DTG rhodopsins are phylogenetically close to DTS rhodopsins, forming a DTG/DTS cluster (Suzuki *et al*. 2022). The DTS rhodopsin from *Sphingomonas paucimobilis* functions as an outward H^+^-pump at pH lower than 7.0 and its photocycle speed is also pH dependent and slows down with increasing extracellular pH (Maliar *et al*. 2020). The pH dependance of DTG/DTS rhodopsins might be due to the lack of a carboxylic H^+^-donating residue as in the DTE/DTD variants (Suzuki *et al*. 2022), and therefore it is plausible that rhodopsins in the DTG/DTS cluster are all pH-dependent. In our study, we found DTG opsin genes in Mycobacteriales and Solirubrobacterales within Actinobacteria, and 121 out of the 130 DTG sequences were from four clearwater lakes with an average pH of 7.6 (averaged from these lakes at depths where these sequences were recovered) (**Figure 2b**). Notably, these four lakes have surface water pH values around 8, which is higher than most other lakes included in the stratfreshDB (Buck *et al*. 2021). By contrast, we found DTS opsin sequences all belonging to Saccharibacteria (SacR) exclusively in humic waters and the average pH of these SacR-occurring waters is 5.2 (**Figure 2b**). Notably, these SacR sequences were initially discovered from the same set of MAGs, functioning as an outward H^+^-pump (Jaffe *et al*. 2022). As characterized DTG and DTS rhodopsins are both pH-dependent outward H^+^ pumps that function more efficiently at low extracellular pH than at high pH, their recovery from non-neutral pH freshwaters in our study is interesting, yet their contrasting occurrence patterns between slightly alkaline and acidic environments seems contradicting. This contradiction may reflect the different aspects of the pH dependance. On one side, stopping pumping out H^+^ at high extracellular pH is beneficial to avoid further cytoplasmic alkalization in slightly alkaline lakes where natural fluctuations of pH can sometimes impose an alkaline pH stress on bacterial cells. On the other side, efficient pumping out H^+^ at low extracellular pH is advantageous for cells to generate energy and/or prevent cytoplasmic acidification in acidic humic waters. This may probably explain the contrasting occurrence patterns of DTG and DTS variants between slightly alkaline clear waters and acidic humic waters and may suggest their association with habitat pH conditions.

In our study, the DTQ motif was exclusively from members of LD12 in humic waters, whereas the typical DTE variant from LD12 occurred in both clear and humic waters. The DTE and DTQ variants were also present in SAR11 (the marine sister clade of LD12), with DTE as the dominant variant in the surface water (0-200 m) (Olson *et al*. 2018). In the same study, a deep-water SAR11 DTQ opsin, when expressed in *Escherichia coli*, functioned as a light-driven outward H^+^-pump, and SAR11 DTQ opsins were expressed especially in deep sea (500–1000 m). This observation, together with the exclusive occurrence of DTQ variants of LD12 in humic waters, may suggest the association of the DTQ variant with poor light conditions.

Unknown variants (i.e. ATT, ATV, or EGV motifs) were exclusively found in humic waters. Interestingly these sequences have the hydrophilic glutamine (Q) residue for spectral tuning (**Figure 2a**) and may represent novel rhodopsins with different functions and/or photochemical behaviors.

As a summary, opsin genes are more widely distributed among clearwater taxa than humic taxa. Conversely, the ion-translocating motif seemed more diverse in humic than in clear waters. The different distributions of these motifs might be associated with the pH, light, or other physicochemical contrasts between clear and humic waters. Further studies are needed to confirm this.

### Significance of rhodopsins in clearwater lakes

Opsin genes are widely distributed in marine environments where they often occur in more than half of the cells in a wide range of samples (Finkel *et al*. 2013; Maresca *et al*. 2018; Olson *et al*. 2018). Some have suggested that opsin genes are more abundant in marine than in freshwater systems (Finkel *et al*. 2013; Oh *et al*. 2011). In particular, opsin gene abundance was positively correlated to salinity (Brindefalk *et al*. 2016; Maresca *et al*. 2018). However, a very limited number of freshwater systems were included in these studies.

We found that in contrast to previous predictions opsin genes are indeed prevalent in clearwater lakes with low DOC and good light penetration. Opsin gene abundance was usually higher than 60% in lakes with DOC less than 5 mg/L. In particular opsin gene abundance is remarkably high (∼90%) in the Laurentian Great Lakes, where DOC is around 2 mg/L and the terrestrial influence is small due to the large surface area. This suggests that rhodopsin-based phototrophy is an important light-harvesting mechanism and its significance may have been underestimated in such lakes. Contrasting to that, opsin genes appear to be less important overall in humic lakes which are often associated with a large terrestrial influence, high allochthonous DOC, brown water color, and a shallow photic zone as discussed in more detail later.

### Opsin gene expression and diel regulation

In Mendota and Sparkling Lake, the majority of bacterial opsin genes were expressed. The total opsin transcripts accounted for 0.3-0.5% of all mRNA, and a few opsin genes were among the most highly expressed genes in these two lakes. These observations support the importance of rhodopsins in bacterial ecophysiology in the surface water of clearwater lakes. Although rhodopsins may also be important in populations that express opsin genes in the bog, the higher total expression of opsin genes in the two clearwater lakes compared to in the humic bog suggests the overall greater significance of rhodopsin-based phototrophy in clearwater lakes.

Opsin gene regulation in freshwater has until now largely unexplored. In marine systems, some studies showed that opsin gene expression exhibited diel rhythms, with maximal expression at the darker periods instead of peak sunlight (e.g. pre-sunrise, post-sunrise, pre-sunset, and post-sunset) (Ottesen *et al*. 2014; Zehnpfennig *et al*. 2022), whereas others suggested opsin genes may be constitutively expressed (Riedel *et al*. 2010; Shi *et al*. 2011). In freshwater, an earlier study of Actinorhodopsin expression indicated their maximal expression at dawn and minimal expression at dusk for samples collected every six hours (Wurzbacher *et al*. 2012). In our study, samples were collected every four hours for two days. From both clearwater lakes, we observed consistent patterns showing diel cyclic behaviors differing among major taxa. Specifically, Actinobacterial opsin genes tend to peak at darker periods of the day, and Bacteroidetes tend to peak during the daytime, especially at peak sunlight.

Taxa-specific diel patterns might suggest differences in a circadian-like clock among bacteria. As opsin gene expression can be regulated by light (Lami *et al*. 2009), carbon availability and nutrient availability (Kwon *et al*. 2013; Wang *et al*. 2012), the taxa-specific patterns might also suggest dual regulation of opsin genes by both light regime and carbon/nutrient availabilities, as the latter can also experience diel cascade changes following photosynthesis in the lake (Linz *et al*. 2020). We should note that only a portion of the expressed opsin genes exhibited a statically significant diel pattern, contributing to about half of all opsin transcripts in the two clearwater lakes (**Figure 6**). It is not clear whether those without cyclic trends are constitutively expressed as some studies suggested (Keffer *et al*. 2015; Riedel *et al*. 2010) or if their diel changes are too small to be detected given the noise in the metatranscriptome data.

### Influence of DOC and optical properties on opsin gene distribution

Lake optical properties are strongly influenced by DOC quality/quantity (Maizel *et al*. 2017; Read & Rose 2013), and in particular, a linear relationship between the light attenuation coefficient for PAR and lake DOC was previously revealed (Read & Rose 2013). Humic lakes often have higher DOC levels than clearwater lakes, as they receive large amounts of terrestrially-derived allochthonous DOC, most of which are in the forms of humic and fulvic acids. These carbon molecules are light-absorbing chromophoric dissolved organic matter (CDOM), mainly absorbing light at wavelengths below 500 nm, including UV-A, UV-B, and the blue light spectrum of photosynthetically active radiation (400-700nm) (Markager & Vincent 2000). In particular, the high concentrations of gilvin and tripton (mostly dissolved and particulate humic substances, respectively) in peat lakes were shown to rapidly absorb blue and green light (Stomp *et al*. 2007). The light spectral distribution can influence phototroph distribution as revealed previously, as increasing humic substance concentration in freshwater increased phototrophs absorbing red light and decreased those absorbing the central part of the PAR spectrum (500–600 nm) (Vila *et al*. 1998). Therefore, as expected, we found that the inferred lower availability of green light strongly limited opsin gene occurrence in humic lakes since the encoded rhodopsins in our analysis were predicted to absorb green light.

Light availability also changes with water depth and such changes differ between clearwater and humic lakes. Opsin gene abundance similarly varied with depth. Depth-associated decreases in opsin gene abundance were relatively moderate and gradual in clearwater lakes. An extreme example is from Lake Michigan, one of the Laurentian Great Lakes with low DOC (∼2 mg/L), where opsin genes remained abundant throughout the water column to 110 m deep. In Lake Tanganyika, the opsin gene abundance estimate was around 30% even at 1100 m. These findings are similar to marine studies, where sunlight penetrates deep into the open ocean, and opsin genes and rhodopsin pigments were abundant in the photic zone (the upper 200 m) (Gomez-Consarnau *et al*. 2019; Olson *et al*. 2018) and opsin genes were still present and expressed at deeper depths (down to 1000 meters, albeit at low abundances) (Olson *et al*. 2018). Therefore, in very large and clear lakes like Michigan and Tanganyika, rhodopsin-based phototrophy may contribute significantly to the total phototrophy and energy budgets within the lake, similar to its counterpart in the open ocean. By contrast, opsin gene abundance quickly declined to very low levels within the first few meters in humic lakes. Therefore, humic lakes not only have lower opsin abundance in the surface layer, but also have a much shallower opsin-containing zone, both contributing to a lower significance of rhodopsin-enabled phototrophy in such environments.

The consistency between light climate and opsin gene profiles with depth suggests the importance of lake optical properties in opsin gene distribution. At the same time, the inverse correlation between opsin gene abundance and DOC levels for samples collected near the surface suggests a strong influence of DOC on opsin gene occurrence. However, as DOC, especially CDOM, and optical properties are generally correlated, it is difficult to tease apart their individual influence on opsin gene distribution from our data. It is plausible that CDOM indirectly influences opsin gene distribution by impacting light availability. In addition, we also postulate a direct influence of DOC on opsin gene distribution, as low DOC may select for opsin-containing bacteria as discussed later.

We should note that DOC not only influences lake optical properties but also influences lake water pH and dissolved oxygen (DO). Humic lakes often have slightly acidic pH (∼5-6, **Table S1**), due to the high levels of humic and fulvic acids. In addition, DO usually decreases sharply along the depth due to lower oxygenic photosynthesis and higher biological oxygen demand as compared to clearwater lakes, leading to a shallow oxygenated zone. Therefore, compared among lakes, acidic pH seems to be associated with low opsin abundance, and compared within a lake, low DO is associated with low opsin abundance. Given that rhodopsin is a light-sensitive protein, the pH and DO influence on opsin gene abundance is likely less important than light availability, although we cannot exclude their potential impacts based on the available data.

### Ecological roles and opsin gene selection in freshwater

Light availability may be one of the main reasons for the wide distribution of opsin genes in clearwater lakes. At the same time, we speculate that competitive advantages for surviving provided by rhodopsins may favor their selection in clearwater lakes. Nearly all bacterial opsin genes that we analyzed encode H^+^-pumps, generating proton motive force (PMF), which contributes to bacterial energy metabolism and renders physiological advantages, especially when bacteria are under stress or carbon-limited conditions as discussed below.

As many vitamins are vulnerable to UV light, vitamins in surface water are subjected to photodegradation in the photic zone. This may impose pressure on microbes to quickly take up vitamins as soon as they are available, particularly in clearwater lakes. Proteorhodopsin gene expression was linked to the light-enhanced growth of two marine vitamin-B_1_ auxotrophic Flavobacteria strains by providing PMF to power the Ton-B dependent acquisition of vitamin-B_1_ (Gomez-Consarnau *et al*. 2016). Therefore, bacteria with H^+^-pumping rhodopsins may compete better for vitamin uptake compared to bacteria without this additional PMF-generating revenue, and are selected for in clearwater lakes where bacteria are subjected to more intense UV radiation than in brown-water humic bogs.

Rhodopsin-containing bacteria are photoheterotrophs relying on organic carbon for growth. Timely carbon substrate scavenging is a key for heterotrophs to succeed under substrate-limited conditions. Due to the additional PMF/ATP generated from light energy, bacteria with H^+^-pumping rhodopsins may be able to enhance the active uptake of organic substrates. More importantly, the energy captured from the sunlight may reduce their reliance on respiration. For example, when organic carbon became limiting, the respiration rate of ‘*Ca.* Pelagibacter ubique’ was lower under light as compared to in the dark while maintaining comparable cell densities (Steindler *et al*. 2011). Therefore, possessing rhodopsins may be more advantageous in carbon-limited than in carbon-rich environments. For example, some rhodopsin-possessing bacteria exhibited light-promoted growth on low organic carbon media or light-enhanced survival under carbon starvation (Gómez-Consarnau *et al*. 2010; Gómez-Consarnau *et al*. 2007; Johnson *et al*. 2010; Steindler *et al*. 2011; Yoshizawa *et al*. 2014). As lakes with high DOC, particularly humic bogs, are not carbon-limited, whereas many clearwater lakes are, rhodopsin-containing bacteria may be strongly selected for in clearwater lakes. This may partly and directly contribute to the inverse correlation between opsin gene abundance and lake DOC levels, in addition to the indirect impact of DOC through its influence on the availability of rhodopsin-absorbing spectra and UV light.

### Rhodopsin-based phototrophy in carbon and energy fluxes in freshwaters

We propose that rhodopsin-based phototrophy influences the carbon and energy fluxes in freshwaters, and the significance of this influence differs between clearwater and humic lakes, as illustrated in **Figure 8**. In clearwater lakes, a large fraction of heterotrophs are actually photoheterotrophs using rhodopsins to harvest sunlight while respiring DOC. As DOC may be limited by photoautotrophic primary producers or present in low levels in clearwater lakes, rhodopsin-phototrophy may be a significant pathway for harvesting energy in such lakes. By contrast, in humic lakes, a large amount of light energy is absorbed by CDOM and particulate humic substances, and such photochemical reactions generate labile DOC, which can be used for heterotrophic respiration (Cory & Kling 2018). As such, although the photodegradation process leaves a small amount of light for rhodopsin-phototrophy, photodegradation provides bacteria with a steady labile organic substrate supply for energy generation. Therefore, compared to clearwater lakes, the contribution of heterotrophy is larger, whereas the contribution of light harvesting by rhodopsins is smaller in the total energy budget in humic lakes. This further influences carbon flux, as the greater reliance on organic carbon based respiration increases the net emission of CO_2_ from humic lakes. It was shown that CO_2_ emission and net heterotrophy were associated with allochthonous DOC concentrations, and were higher in humic than in clearwater lakes (Jonsson *et al*. 2003). From our study, it is possible that rhodopsin-based phototrophy may be a previously overlooked factor contributing to such a difference in CO_2_ emission from these two lake types.

**Figure 8.**
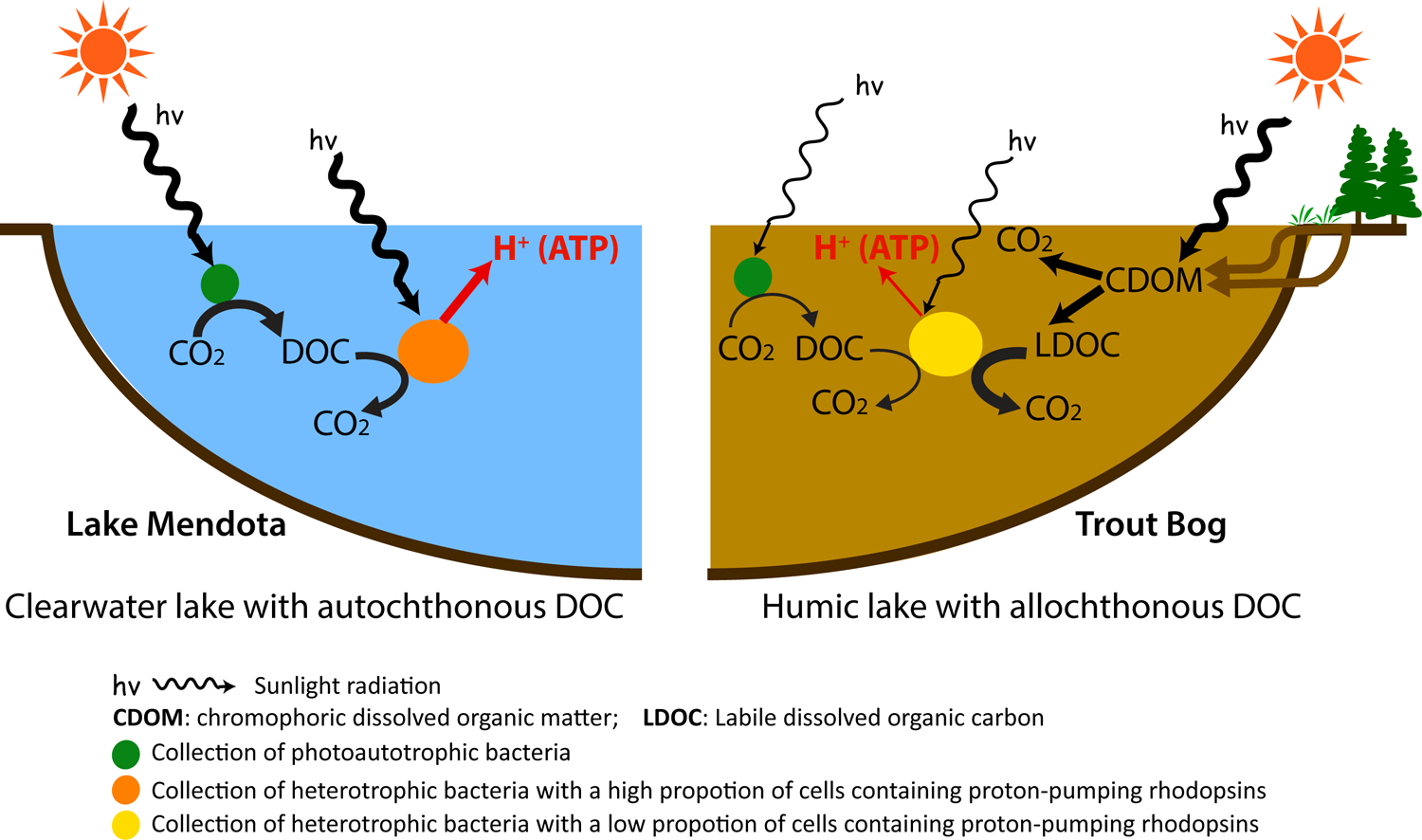
Difference on the significance of rhodopsin-based phototrophy in carbon metabolism and energy flux between clearwater lakes (such as Lake Mendota) and humic lakes (such as Trout Bog). Prevalent of reactions is qualitatively indicated by the thickness of lines.

## SUMMARY

In this study, by analyzing opsin genes recovered in metagenomes and MAGs collected from clear and humic waters with contrasting pH, optical properties and DOC quantity/quality, we revealed differences in opsin diversity between two major freshwater types and recovered novel diversity within freshwater opsin genes. Many of these opsin genes were expressed, and some of them exhibited diel cyclic rhythms that differ among taxa. Further, we found opsin gene abundance is associated with lake DOC and depth in the water column, suggesting their ecological distribution can be influenced by organic matter and lake optical properties. In particular, opsin genes are widely present and highly expressed in clearwater lakes, and may be strongly selected for due to the higher light availability, lower DOC, and thus greater competitive advantages provided by rhodopsins in such environments. Therefore, we assert that rhodopsin-based phototrophy is a significant light harvesting mechanism, substantially influencing energy and carbon budgets in clearwater lakes.

## ACKNOWLEDGEMENTS

We thank the North Temperate Lakes Microbial Observatory 2007-2012 field crews, UW-Trout Lake Station, the UW Center for Limnology, and the Global Lakes Ecological Observatory Network for field and logistical support. We acknowledge efforts by many McMahon Lab undergrads and technicians related to sample collection and DNA extraction. We thank the individual program directors and leadership at the National Science Foundation for their commitment to the continued support of long-term ecological research. We also thank Dr. Andrey Rozenberg for providing valuable and helpful comments on the preprint of this manuscript. The project was supported by funding from the United States National Science Foundation Microbial Observatories program (MCB-0702395), the Long-Term Ecological Research program (NTL-LTER DEB-1440297), an INSPIRE award (DEB-1344254) and U.S. Department of Agriculture (Hatch Project WIS03052). This material is also based upon work supported by the National Institute of Food and Agriculture, U.S. Department of Agriculture (Hatch Project 1002996). The work conducted by the U.S. Department of Energy Joint Genome Institute, a DOE Office of Science User Facility, is supported by the Office of Science of the U.S. Department of Energy under Contract No. DE-AC02-05CH11231.

## Data Accessibility and Benefit-Sharing

All metagenome data are publicly available at the DOE Joint Genome Institute’s Integrated Microbial Genome (IMG) database or European Nucleotide Archive, with their Accession No. listed in Tables S2, S3, and S4. All metatranscriptome data are from the study by Linz et al. (2020). Amino acid sequences of opsin genes used in this study are listed in Supplementary Files 1, 2, 3, and 4.

## Author Contributions

K.D.M and S.H. designed and initiated the study; A.M.L. generated the metatranscriptome and part of the metagenome data; S.S. created the metagenome bins; P.T. generated part of the metagenome data; F.M.-F., B.O.O., J.R.D.-S., K.T.F., K.D.M., S.H., A.M.L. and S.S. participated in study discussion and metatranscriptome sampling; S.H. performed data analysis and interpreted the results; S.H. and K.D.M. wrote the manuscript. All authors contributed to the discussion and improvement of the manuscript.

## REFERENCES

1. Atamna-Ismaeel N, Sabehi G, Sharon I, et al. (2008) Widespread distribution of proteorhodopsins in freshwater and brackish ecosystems. ISME J 2, 656–662.

2. Balashov SP, Imasheva ES, Boichenko VA, et al. (2005) Xanthorhodopsin: A Proton Pump with a Light-Harvesting Carotenoid Antenna. Science 309, 2061–2064.

3. Balashov SP, Petrovskaya LE, Imasheva ES, et al. (2013) Breaking the carboxyl rule: lysine 96 facilitates reprotonation of the Schiff base in the photocycle of a retinal protein from Exiguobacterium sibiricum. J Biol Chem 288, 21254–21265.

4. Béjà O, Aravind L, Koonin EV, et al. (2000) Bacterial Rhodopsin: Evidence for a New Type of Phototrophy in the Sea. Science 289, 1902–1906.

5. Beja O, Lanyi JK (2014) Nature’s toolkit for microbial rhodopsin ion pumps. Proc Natl Acad Sci U S A 111, 6538–6539.

6. Bendall ML, Stevens SLR, Chan L-K, et al. (2016) Genome-wide selective sweeps and gene-specific sweeps in natural bacterial populations. ISME J.

7. Bratanov D, Kovalev K, Machtens JP, et al. (2019) Unique structure and function of viral rhodopsins. Nature Communications 10.

8. Brindefalk B, Ekman M, Ininbergs K, et al. (2016) Distribution and expression of microbial rhodopsins in the Baltic Sea and adjacent waters. Environ Microbiol 18, 4442–4455.

9. Buck M, Garcia SL, Fernandez L, et al. (2021) Comprehensive dataset of shotgun metagenomes from oxygen stratified freshwater lakes and ponds. Scientific Data 8.

10. Cabello-Yeves PJ, Ghai R, Mehrshad M, et al. (2017) Reconstruction of Diverse Verrucomicrobial Genomes from Metagenome Datasets of Freshwater Reservoirs. Frontiers in Microbiology 8.

11. Castresana J (2000) Selection of conserved blocks from multiple alignments for their use in phylogenetic analysis. Mol Biol Evol 17.

12. Chazan A, Rozenberg A, Mannen K, et al. (2022) Diverse heliorhodopsins detected via functional metagenomics in freshwater Actinobacteria, Chloroflexi and Archaea. Environ Microbiol 24, 110–121.

13. Cory RM, Kling GW (2018) Interactions between sunlight and microorganisms influence dissolved organic matter degradation along the aquatic continuum. Limnology and Oceanography Letters 3, 102–116.

14. Dam P, Rodriguez-R LM, Luo CW, et al. (2020) Model-based Comparisons of the Abundance Dynamics of Bacterial Communities in Two Lakes. Scientific Reports 10.

15. Darling AE, Jospin G, Lowe E, et al. (2014) PhyloSift: phylogenetic analysis of genomes and metagenomes. PeerJ 2, e243.

16. Denef VJ, Mueller RS, Chiang E, Liebig JR, Vanderploeg HA (2016) Chloroflexi CL500-11 Populations That Predominate Deep-Lake Hypolimnion Bacterioplankton Rely on Nitrogen-Rich Dissolved Organic Matter Metabolism and C(1) Compound Oxidation. Applied and Environmental Microbiology 82, 1423–1432.

17. Durán-Viseras A, Andrei A-S, Ghai R, Sánchez-Porro C, Ventosa A (2019) New Halonotius Species Provide Genomics-Based Insights Into Cobalamin Synthesis in Haloarchaea. Frontiers in Microbiology 10.

18. Dwulit-Smith JR, Hamilton JJ, Stevenson DM, et al. (2018) acI Actinobacteria Assemble a Functional Actinorhodopsin with Natively Synthesized Retinal. Applied and Environmental Microbiology 84.

19. Edgar RC (2004) MUSCLE: a multiple sequence alignment method with reduced time and space complexity. BMC Bioinformatics 5, 113–113.

20. Edgar RC (2010) Search and clustering orders of magnitude faster than BLAST. Bioinformatics 26, 2460–2461.

21. Finkel OM, Beja O, Belkin S (2013) Global abundance of microbial rhodopsins. ISME J 7, 448–451.

22. Flores-Uribe J, Hevroni G, Ghai R, et al. (2019) Heliorhodopsins are absent in diderm (Gram-negative) bacteria: Some thoughts and possible implications for activity. Environ Microbiol Rep 11, 419–424.

23. Galachyants AD, Krasnopeev AY, Podlesnaya GV, et al. (2021) Diversity of Aerobic Anoxygenic Phototrophs and Rhodopsin-Containing Bacteria in the Surface Microlayer, Water Column and Epilithic Biofilms of Lake Baikal. Microorganisms 9.

24. Garcia SL, McMahon KD, Grossart H-P, Warnecke F (2014) Successful enrichment of the ubiquitous freshwater acI Actinobacteria. Environ Microbiol Rep 6, 21–27.

25. Ghai R, Mizuno CM, Picazo A, Camacho A, Rodriguez-Valera F (2014) Key roles for freshwater Actinobacteria revealed by deep metagenomic sequencing. Molecular Ecology 23, 6073–6090.

26. Gifford SM, Sharma S, Rinta-Kanto JM, Moran MA (2011) Quantitative analysis of a deeply sequenced marine microbial metatranscriptome. Isme Journal 5, 461–472.

27. Gómez-Consarnau L, Akram N, Lindell K, et al. (2010) Proteorhodopsin Phototrophy Promotes Survival of Marine Bacteria during Starvation. PLoS Biology 8, e1000358.

28. Gómez-Consarnau L, Gonzalez JM, Coll-Llado M, et al. (2007) Light stimulates growth of proteorhodopsin-containing marine Flavobacteria. Nature 445, 210–213.

29. Gomez-Consarnau L, Gonzalez JM, Riedel T, et al. (2016) Proteorhodopsin light-enhanced growth linked to vitamin-B1 acquisition in marine Flavobacteria. ISME J 10, 1102–1112.

30. Gomez-Consarnau L, Raven JA, Levine NM, et al. (2019) Microbial rhodopsins are major contributors to the solar energy captured in the sea. Science Advances 5.

31. Guindon S, Dufayard JF, Lefort V, et al. (2010) New algorithms and methods to estimate maximum-likelihood phylogenies: assessing the performance of PhyML 3.0. Syst Biol 59, 307–321.

32. Harris A, Ljumovic M, Bondar AN, et al. (2015) A new group of eubacterial light-driven retinal-binding proton pumps with an unusual cytoplasmic proton donor. Biochim Biophys Acta 1847, 1518–1529.

33. He S, Malfatti SA, McFarland JW, et al. (2015) Patterns in wetland microbial community composition and functional gene repertoire associated with methane emissions. mBio 6, e00066–00015.

34. Imasheva ES, Balashov SP, Choi AR, Jung K-H, Lanyi JK (2009) Reconstitution of Gloeobacter violaceus Rhodopsin with a Light-Harvesting Carotenoid Antenna. Biochemistry 48, 10948–10955.

35. Jaffe AL, Konno M, Kawasaki Y, et al. (2022) Saccharibacteria harness light energy using type-1 rhodopsins that may rely on retinal sourced from microbial hosts. The ISME Journal 16, 2056–2059.

36. Johnson ET, Baron DB, Naranjo B, et al. (2010) Enhancement of survival and electricity production in an engineered bacterium by light-driven proton pumping. Appl Environ Microbiol 76, 4123–4129.

37. Jonsson A, Karlsson J, Jansson M (2003) Sources of Carbon Dioxide Supersaturation in Clearwater and Humic Lakes in Northern Sweden. Ecosystems 6, 224–235.

38. Keffer JL, Hahn MW, Maresca JA (2015) Characterization of an Unconventional Rhodopsin from the Freshwater Actinobacterium Rhodoluna lacicola. J Bacteriol 197, 2704–2712.

39. Kwon SK, Kim BK, Song JY, et al. (2013) Genomic makeup of the marine flavobacterium Nonlabens (Donghaeana) dokdonensis and identification of a novel class of rhodopsins. Genome Biol Evol 5, 187–199.

40. Lami R, Cottrell MT, Campbell BJ, Kirchman DL (2009) Light-dependent growth and proteorhodopsin expression by Flavobacteria and SAR11 in experiments with Delaware coastal waters. Environ Microbiol 11, 3201–3209.

41. Le SQ, Gascuel O (2008) An improved general amino acid replacement matrix. Mol Biol Evol 25, 1307–1320.

42. Linz AM, Aylward FO, Bertilsson S, McMahon KD (2020) Time-series metatranscriptomes reveal conserved patterns between phototrophic and heterotrophic microbes in diverse freshwater systems. Limnology and Oceanography 65, S101–S112.

43. Linz AM, He SM, Stevens SLR, et al. (2018) Freshwater carbon and nutrient cycles revealed through reconstructed population genomes. Peerj 6.

44. Maizel AC, Li J, Remucal CK (2017) Relationships Between Dissolved Organic Matter Composition and Photochemistry in Lakes of Diverse Trophic Status. Environmental Science & Technology 51, 9624–9632.

45. Maliar N, Okhrimenko IS, Petrovskaya LE, et al. (2020) Novel pH-Sensitive Microbial Rhodopsin from Sphingomonas paucimobilis. Doklady Biochemistry and Biophysics 495, 342–346.

46. Man D, Wang W, Sabehi G, et al. (2003) Diversification and spectral tuning in marine proteorhodopsins. The EMBO Journal 22, 1725–1731.

47. Maresca JA, Miller KJ, Keffer JL, Sabanayagam CR, Campbell BJ (2018) Distribution and Diversity of Rhodopsin-Producing Microbes in the Chesapeake Bay. Applied and Environmental Microbiology 84.

48. Markager S, Vincent WF (2000) Spectral light attenuation and the absorption of UV and blue light in natural waters. Limnology and Oceanography 45, 642–650.

49. Markowitz VM, Chen I-MA, Palaniappan K, et al. (2013) IMG 4 version of the integrated microbial genomes comparative analysis system. Nucleic Acids Research.

50. Martinez-Garcia M, Swan BK, Poulton NJ, et al. (2012) High-throughput single-cell sequencing identifies photoheterotrophs and chemoautotrophs in freshwater bacterioplankton. ISME J 6.

51. Matsen FA, Kodner RB, Armbrust EV (2010) pplacer: linear time maximum-likelihood and Bayesian phylogenetic placement of sequences onto a fixed reference tree. BMC Bioinformatics 11, 538.

52. McCarren J, DeLong EF (2007) Proteorhodopsin photosystem gene clusters exhibit co-evolutionary trends and shared ancestry among diverse marine microbial phyla. Environ Microbiol 9, 846–858.

53. Mizuno CM, Rodriguez-Valera F, Ghai R (2015) Genomes of Planktonic Acidimicrobiales: Widening Horizons for Marine Actinobacteria by Metagenomics. mBio 6.

54. Needham DM, Yoshizawa S, Hosaka T, et al. (2019) A distinct lineage of giant viruses brings a rhodopsin photosystem to unicellular marine predators. Proc Natl Acad Sci U S A 116, 20574–20583.

55. Newton RJ, Kent AD, Triplett EW, McMahon KD (2006) Microbial community dynamics in a humic lake: differential persistence of common freshwater phylotypes. Environ Microbiol 8, 956–970.

56. Oh S, Caro-Quintero A, Tsementzi D, et al. (2011) Metagenomic Insights into the Evolution, Function, and Complexity of the Planktonic Microbial Community of Lake Lanier, a Temperate Freshwater Ecosystem. Applied and Environmental Microbiology 77, 6000–6011.

57. Olson DK, Yoshizawa S, Boeuf D, Iwasaki W, DeLong EF (2018) Proteorhodopsin variability and distribution in the North Pacific Subtropical Gyre. Isme Journal 12, 1047–1060.

58. Ottesen EA, Young CR, Gifford SM, et al. (2014) Multispecies diel transcriptional oscillations in open ocean heterotrophic bacterial assemblages. Science 345, 207–212.

59. Philosof A, Beja O (2013) Bacterial, archaeal and viral-like rhodopsins from the Red Sea. Environ Microbiol Rep 5, 475–482.

60. Pinhassi J, DeLong EF, Beja O, Gonzalez JM, Pedros-Alio C (2016) Marine Bacterial and Archaeal Ion-Pumping Rhodopsins: Genetic Diversity, Physiology, and Ecology. Microbiology and Molecular Biology Reviews 80.

61. Podowski JC, Paver SF, Newton RJ, Coleman ML (2021) Genome streamlining, proteorhodopsin, and organic nitrogen metabolism in freshwater nitrifiers. bioRxiv, 2021.2001.2019.427344.

62. Pushkarev A, Inoue K, Larom S, et al. (2018) A distinct abundant group of microbial rhodopsins discovered using functional metagenomics. Nature 558, 595-+.

63. Read JS, Rose KC (2013) Physical responses of small temperate lakes to variation in dissolved organic carbon concentrations. Limnology and Oceanography 58, 921–931.

64. Riedel T, Tomasch J, Buchholz I, et al. (2010) Constitutive Expression of the Proteorhodopsin Gene by a Flavobacterium Strain Representative of the Proteorhodopsin-Producing Microbial Community in the North Sea. Applied and Environmental Microbiology 76, 3187–3197.

65. Rozenberg A, Inoue K, Kandori H, Beja O (2021) Microbial Rhodopsins: The Last Two Decades. In: Annual Review of Microbiology, Vol 75, 2021 (ed. Gottesman S), pp. 427-447.

66. Rusch DB, Halpern AL, Sutton G, et al. (2007) The Sorcerer II Global Ocean Sampling expedition: northwest Atlantic through eastern tropical Pacific. PLoS Biol 5.

67. Sharma AK, Sommerfeld K, Bullerjahn GS, et al. (2009) Actinorhodopsin genes discovered in diverse freshwater habitats and among cultivated freshwater Actinobacteria. ISME J 3, 726–737.

68. Sharma AK, Zhaxybayeva O, Papke RT, Doolittle WF (2008) Actinorhodopsins: proteorhodopsin-like gene sequences found predominantly in non-marine environments. Environ Microbiol 10, 1039–1056.

69. Shi YM, Tyson GW, Eppley JM, DeLong EF (2011) Integrated metatranscriptomic and metagenomic analyses of stratified microbial assemblages in the open ocean. Isme Journal 5, 999–1013.

70. Sieradzki ET, Fuhrman JA, Rivero-Calle S, Gomez-Consamau L (2018) Proteorhodopsins dominate the expression of phototrophic mechanisms in seasonal and dynamic marine picoplankton communities. Peerj 6.

71. Steindler L, Schwalbach MS, Smith DP, Chan F, Giovannoni SJ (2011) Energy starved Candidatus Pelagibacter ubique substitutes light-mediated ATP production for endogenous carbon respiration. PLoS One 6, e19725.

72. Stomp M, Huisman J, Stal LJ, Matthijs HC (2007) Colorful niches of phototrophic microorganisms shaped by vibrations of the water molecule. ISME J 1, 271–282.

73. Sudo Y, Yoshizawa S (2016) Functional and Photochemical Characterization of a Light-Driven Proton Pump from the Gammaproteobacterium Pantoea vagans. Photochem Photobiol 92, 420–427.

74. Suzuki K, Del Carmen Marín M, Konno M, et al. (2022) Structural characterization of proton-pumping rhodopsin lacking a cytoplasmic proton donor residue by X-ray crystallography. J Biol Chem 298, 101722.

75. Thaben PF, Westermark PO (2014) Detecting Rhythms in Time Series with RAIN. Journal of Biological Rhythms 29, 391–400.

76. Tran P, Ramachandran A, Khawasik O, et al. (2018) Microbial life under ice: Metagenome diversity and in situ activity of Verrucomicrobia in seasonally ice-covered Lakes. Environ Microbiol 20, 2568–2584.

77. Tran PQ, Bachand SC, McIntyre PB, et al. (2021) Depth-discrete metagenomics reveals the roles of microbes in biogeochemical cycling in the tropical freshwater Lake Tanganyika. ISME J 15, 1971–1986.

78. Vila X, Cristina XP, Abella CA, Hurley JP (1998) Effects of gilvin on the composition and dynamics of metalimnetic communities of phototrophic bacteria in freshwater North-American lakes. J Appl Microbiol 85 Suppl 1, 138S–150S.

79. Vollmers J, Voget S, Dietrich S, et al. (2013) Poles apart: Arctic and Antarctic Octadecabacter strains share high genome plasticity and a new type of xanthorhodopsin. PLoS One 8, e63422.

80. Wang Z, O’Shaughnessy TJ, Soto CM, et al. (2012) Function and Regulation of Vibrio campbellii Proteorhodopsin: Acquired Phototrophy in a Classical Organoheterotroph. PLoS One 7.

81. Wurzbacher C, Salka I, Grossart HP (2012) Environmental actinorhodopsin expression revealed by a new in situ filtration and fixation sampler. Environ Microbiol Rep 4, 491–497.

82. Yoshizawa S, Kumagai Y, Kim H, et al. (2014) Functional characterization of flavobacteria rhodopsins reveals a unique class of light-driven chloride pump in bacteria. Proc Natl Acad Sci U S A 111, 6732–6737.

83. Yutin N, Koonin EV (2012) Proteorhodopsin genes in giant viruses. Biology Direct 7, 34–34.

84. Zabelskii D, Alekseev A, Kovalev K, et al. (2020) Viral rhodopsins 1 are an unique family of light-gated cation channels. Nature Communications 11.

85. Zehnpfennig JR, Hansel CM, Wankel SD, et al. (2022) Diel Patterns in Marine Microbial Metatranscriptomes Reflect Differences in Community Metabolic Activity Over Depth on the Continental Shelf of the North Atlantic. Frontiers in Marine Science 9.

