## Supplemental Figure 1 for "Diversity, distribution, and expression of opsin genes in freshwater lakes"

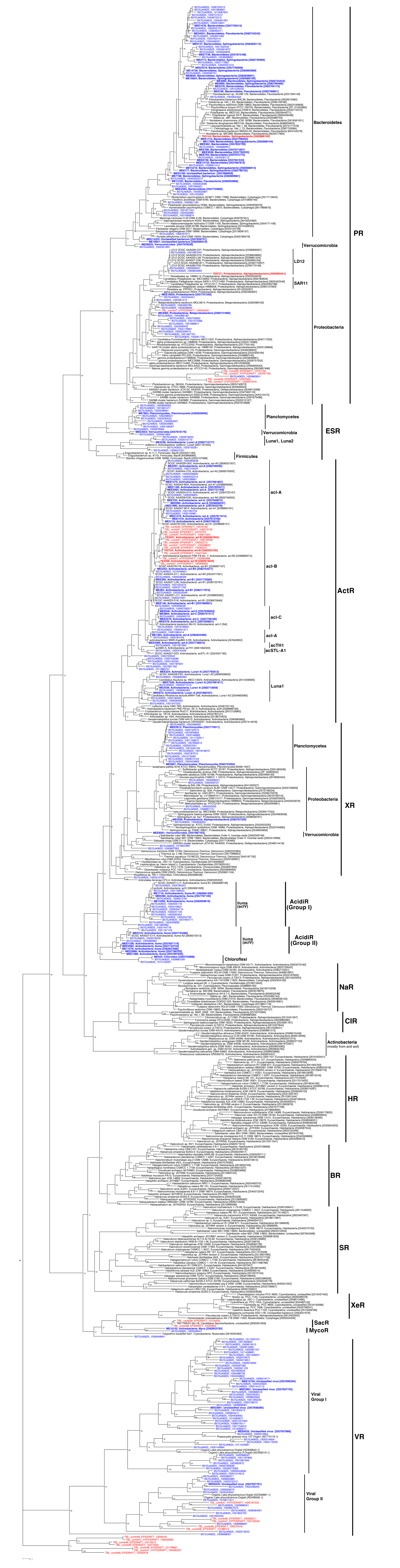
PR
Bacteroidetes
Verrucomicrobia
LD12
SAR11
Proteobacteria
Planctomycetes
Verrucomicrobia
Luna1, Luna2
ESR
Firmicutes
acI-A
acI-B
acI-C
acI-A
acTH1
acSTL-A1
ActR
Luna1
Planctomycetes
Proteobacteria
Verrucomicrobia
XR
Iluma (acIV)
AcidiR (Group I)
Iluma (acIV)
AcidiR (Group II)
Chloroflexi
NaR
ClR
Actinobacteria (mostly from soil)
HR
BR
SR
XeR
SacR
MycoR
Viral Group I
VR
Viral Group II

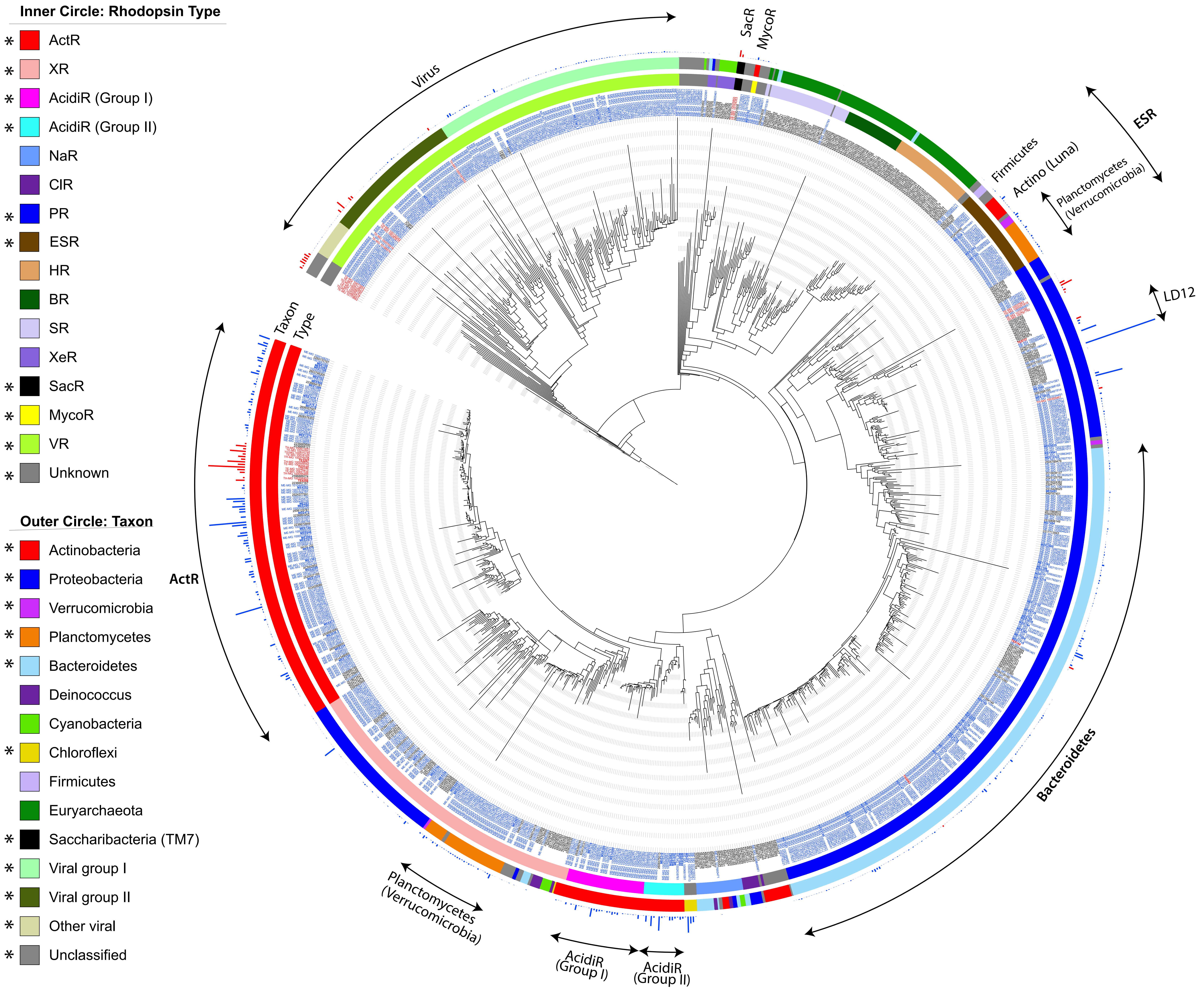
Inner Circle: Rhodopsin Type
* ActR
* XR
* AcidiR (Group I)
* AcidiR (Group II)
NaR
ClR
* PR
* ESR
HR
BR
SR
XeR
* SacR
* MycoR
* VR
* Unknown
Outer Circle: Taxon
* Actinobacteria
* Proteobacteria
* Verrucomicrobia
* Planctomycetes
* Bacteroidetes
Deinococcus
Cyanobacteria
* Chloroflexi
Firmicutes
Euryarchaeota
* Saccharibacteria (TM7)
* Viral group I
* Viral group II
* Other viral
* Unclassified
Virus
SacR
MycoR
ESR
Firmicutes
Actino (Luna)
Planctomycetes (Verrucomicrobia)
LD12
Taxon
Type
ActR
Bacteroidetes
Planctomycetes (Verrucomicrobia)
AcidiR (Group I)
AcidiR (Group II)

### Breakdown by taxa

| | Relative abundance among all opsin genes (%) | Actinobacteria | | | | | | | | | Bacteroidetes | Proteobacteria | Planctomycetes | Verrucomicrobia | Chloroflexi | Sacchribacteria | Virus | | Unclassified |
|---|---|---|---|---|---|---|---|---|---|---|---|---|---|---|---|---|---|---|---|
| | | acI-A | acI-B | acI-C | acTH1 | acSTL | Luna | Myco | Iluma | Unclassified | | | | | | | Group I | Group II | |
| **ME** | | | | | | | | | | | | | | | | | | | |
| ActR | 40.6 | 12.5 | 16.3 | 3.2 | 0.1 | 0.7 | 3.9 | | | 4.0 | | | | | | | | | |
| AcidiR (Group I) | 4.8 | | | | | | | | 4.8 | | | | | | | | | | |
| AcidiR (Group II) | 5.7 | | | | | | | | 5.7 | | | | | | | | | | |
| XR | 6.5 | | | | | | | | | 0.02 | | 2.7 | 3.4 | 0.1 | | | | | 0.2 |
| PR | 28.8 | | | | | | | | | | 12.6 | 16.0 | | 0.1 | | | | | 0.2 |
| ESR | 3.0 | | | | | | 0.7 | | | | | | 1.5 | 0.8 | | | | | |
| MycoR | 0.3 | | | | | | | 0.3 | | | | | | | | | | | |
| VR | 6.0 | | | | | | | | | | | | | | | | 5.0 | 1.1 | |
| Unknown | 4.0 | | | | | | | | | 0.1 | | | | | 3.4 | | | | 0.4 |
| Sum by phylum | | **52.3** | | | | | | | | | **12.6** | **18.7** | **4.9** | **1.0** | **3.4** | **0.0** | **6.0** | | **0.8** |
| **TE** | | | | | | | | | | | | | | | | | | | |
| ActR | 61.3 | | 61.3 | | | | | | | | | | | | | | | | |
| PR | 10.4 | | | | | | | | | | 2.9 | 7.5 | | | | | | | |
| SacR | 5.4 | | | | | | | | | | | | | | | 5.4 | | | |
| VR | 9.9 | | | | | | | | | | | | | | | | | 9.9 | |
| Unknown | 12.9 | | | | | | | | | | | | | | | | | | 12.9 |
| Sum by phylum | | **61.3** | | | | | | | | | **2.9** | **7.5** | | | | **5.4** | **9.9** | | **12.9** |
| **TH** | | | | | | | | | | | | | | | | | | | |
| ActR | 46.3 | | 46.3 | | | | | | | | | | | | | | | | |
| PR | 33.4 | | | | | | | | | | 2.5 | 30.9 | | | | | | | |
| VR | 10.2 | | | | | | | | | | | | | | | | | 10.2 | |
| Unknown | 10.1 | | | | | | | | | | | | | | | | | | 10.1 |
| Sum by phylum | | **46.3** | | | | | | | | | **2.5** | **30.9** | | | | | **10.2** | | **10.1** |

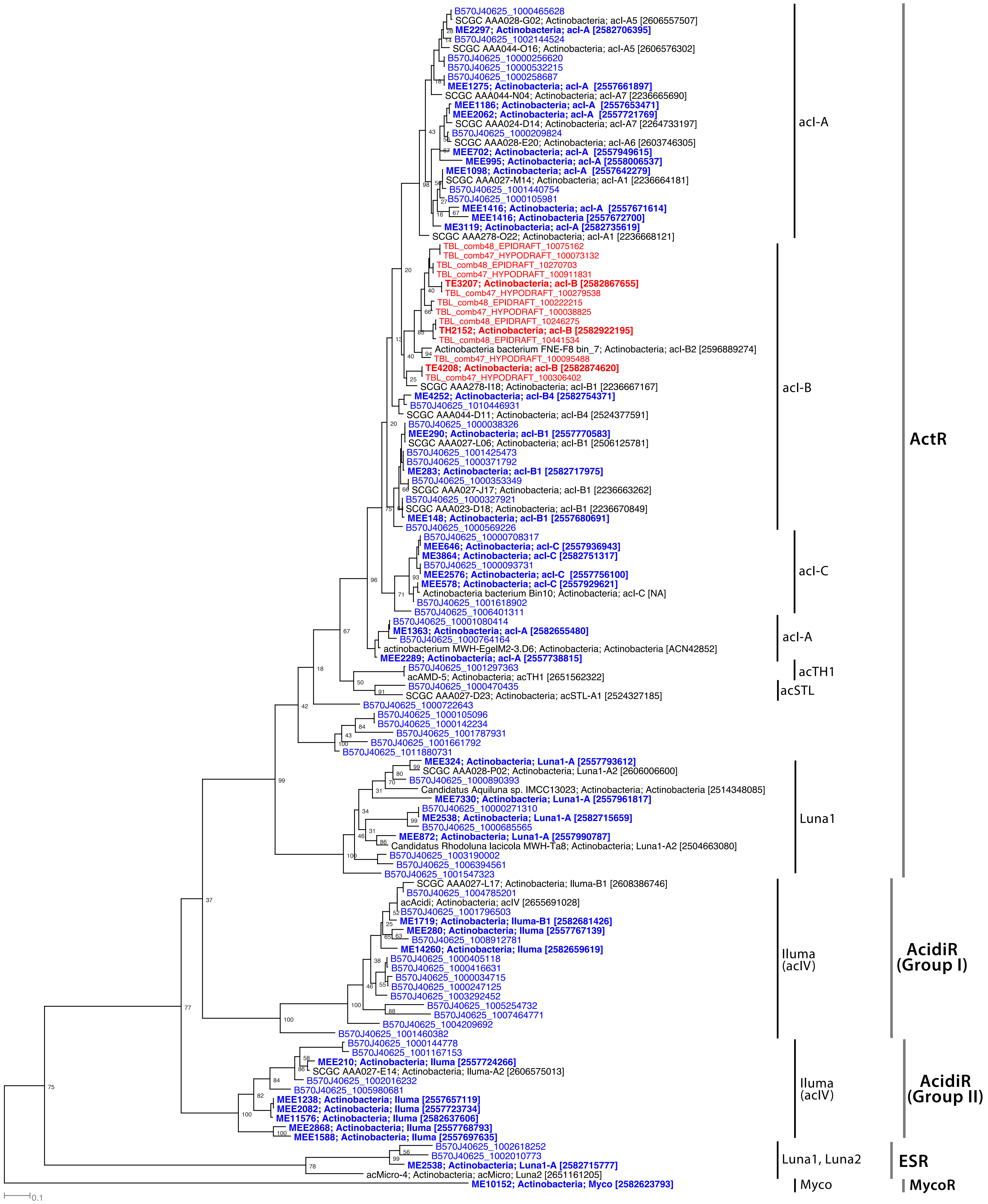

B570J40625_1000465628
SCGC AAA028-G02; Actinobacteria; acI-A5 [2606557507]
ME2297; Actinobacteria; acI-A [2582706395]
B570J40625_1002144524
SCGC AAA044-O16; Actinobacteria; acI-A5 [2606576302]
B570J40625_10000256620
B570J40625_10000532215
B570J40625_1000258687
MEE1275; Actinobacteria; acI-A [2557661897]
SCGC AAA044-N04; Actinobacteria; acI-A7 [2236665690]
MEE1186; Actinobacteria; acI-A [2557653471]
MEE2062; Actinobacteria; acI-A [2557721769]
SCGC AAA024-D14; Actinobacteria; acI-A7 [2264733197]
B570J40625_1000209824
SCGC AAA028-E20; Actinobacteria; acI-A6 [2603746305]
MEE702; Actinobacteria; acI-A [2557949615]
MEE995; Actinobacteria; acI-A [2558006537]
MEE1098; Actinobacteria; acI-A [2557642279]
SCGC AAA027-M14; Actinobacteria; acI-A1 [2236664181]
B570J40625_1001440754
B570J40625_1000105981
MEE1416; Actinobacteria; acI-A [2557671614]
MEE1416; Actinobacteria [2557672700]
ME3119; Actinobacteria; acI-A [2582735619]
SCGC AAA278-O22; Actinobacteria; acI-A1 [2236668121]
TBL_comb48_EPIDRAFT_10075162
TBL_comb47_HYPODRAFT_100073132
TBL_comb48_EPIDRAFT_10270703
TBL_comb47_HYPODRAFT_100911831
TE3207; Actinobacteria; acI-B [2582867655]
TBL_comb47_HYPODRAFT_100279538
TBL_comb48_EPIDRAFT_100222215
TBL_comb47_HYPODRAFT_100038825
TBL_comb48_EPIDRAFT_10246275
TH2152; Actinobacteria; acI-B [2582922195]
TBL_comb48_EPIDRAFT_10441534
Actinobacteria bacterium FNE-F8 bin_7; Actinobacteria; acI-B2 [2596889274]
TBL_comb47_HYPODRAFT_100095488
TE4208; Actinobacteria; acI-B [2582874620]
TBL_comb47_HYPODRAFT_100306402
SCGC AAA278-I18; Actinobacteria; acI-B1 [2236667167]
ME4252; Actinobacteria; acI-B4 [2582754371]
B570J40625_1010446931
SCGC AAA044-D11; Actinobacteria; acI-B4 [2524377591]
B570J40625_1000038326
MEE290; Actinobacteria; acI-B1 [2557770583]
SCGC AAA027-L06; Actinobacteria; acI-B1 [2506125781]
B570J40625_1001425473
B570J40625_1000371792
ME283; Actinobacteria; acI-B1 [2582717975]
B570J40625_1000353349
SCGC AAA027-J17; Actinobacteria; acI-B1 [2236663262]
B570J40625_1000327921
SCGC AAA023-D18; Actinobacteria; acI-B1 [2236670849]
MEE148; Actinobacteria; acI-B1 [2557680691]
B570J40625_1000569226
B570J40625_10000708317
MEE646; Actinobacteria; acI-C [2557936943]
ME3864; Actinobacteria; acI-C [2582751317]
B570J40625_1000093731
MEE2576; Actinobacteria; acI-C [2557756100]
MEE578; Actinobacteria; acI-C [2557929621]
Actinobacteria bacterium Bin10; Actinobacteria; acI-C [NA]
B570J40625_1001618902
B570J40625_1006401311
B570J40625_10001080414
ME1363; Actinobacteria; acI-A [2582655480]
B570J40625_1000764164
actinobacterium MWH-EgelM2-3.D6; Actinobacteria; Actinobacteria [ACN42852]
MEE2289; Actinobacteria; acI-A [2557738815]
B570J40625_1001297363
acAMD-5; Actinobacteria; acTH1 [2651562322]
B570J40625_1000470435
SCGC AAA027-D23; Actinobacteria; acSTL-A1 [2524327185]
B570J40625_1000722643
B570J40625_1000105096
B570J40625_1000142234
B570J40625_1001787931
B570J40625_1001661792
B570J40625_1011880731
MEE324; Actinobacteria; Luna1-A [2557793612]
SCGC AAA028-P02; Actinobacteria; Luna1-A2 [2606006600]
B570J40625_1000890393
Candidatus Aquiluna sp. IMCC13023; Actinobacteria; Actinobacteria [2514348085]
MEE7330; Actinobacteria; Luna1-A [2557961817]
B570J40625_1000271310
ME2538; Actinobacteria; Luna1-A [2582715659]
B570J40625_1000685565
MEE872; Actinobacteria; Luna1-A [2557990787]
Candidatus Rhodoluna lacicola MWH-Ta8; Actinobacteria; Luna1-A2 [2504663080]
B570J40625_1003190002
B570J40625_1006394561
B570J40625_1001547323
SCGC AAA027-L17; Actinobacteria; Iluma-B1 [2608386746]
B570J40625_1004785201
acAcidi; Actinobacteria; acIV [2655691028]
B570J40625_1001796503
ME1719; Actinobacteria; Iluma-B1 [2582681426]
MEE280; Actinobacteria; Iluma [2557767139]
B570J40625_1008912781
ME14260; Actinobacteria; Iluma [2582659619]
B570J40625_1000405118
B570J40625_1000416631
B570J40625_1000034715
B570J40625_1000247125
B570J40625_1003292452
B570J40625_1005254732
B570J40625_1007464771
B570J40625_1004209692
B570J40625_1001460382
B570J40625_1000144778
B570J40625_1001167153
MEE210; Actinobacteria; Iluma [2557724266]
SCGC AAA027-E14; Actinobacteria; Iluma-A2 [2606575013]
B570J40625_1002016232
B570J40625_1005980681
MEE1238; Actinobacteria; Iluma [2557657119]
MEE2082; Actinobacteria; Iluma [2557723734]
ME11576; Actinobacteria; Iluma [2582637606]
MEE2868; Actinobacteria; Iluma [2557768793]
MEE1588; Actinobacteria; Iluma [2557697635]
B570J40625_1002618252
B570J40625_1002010773
ME2538; Actinobacteria; Luna1-A [2582715777]
acMicro-4; Actinobacteria; acMicro; Luna2 [2651161205]
ME10152; Actinobacteria; Myco [2582623793]
0.1
acI-A
acI-B
acI-C
acI-A
acTH1
acSTL
Luna1
Iluma (acIV)
Iluma (acIV)
Luna1, Luna2
Myco
ActR
AcidiR (Group I)
AcidiR (Group II)
ESR
MycoR

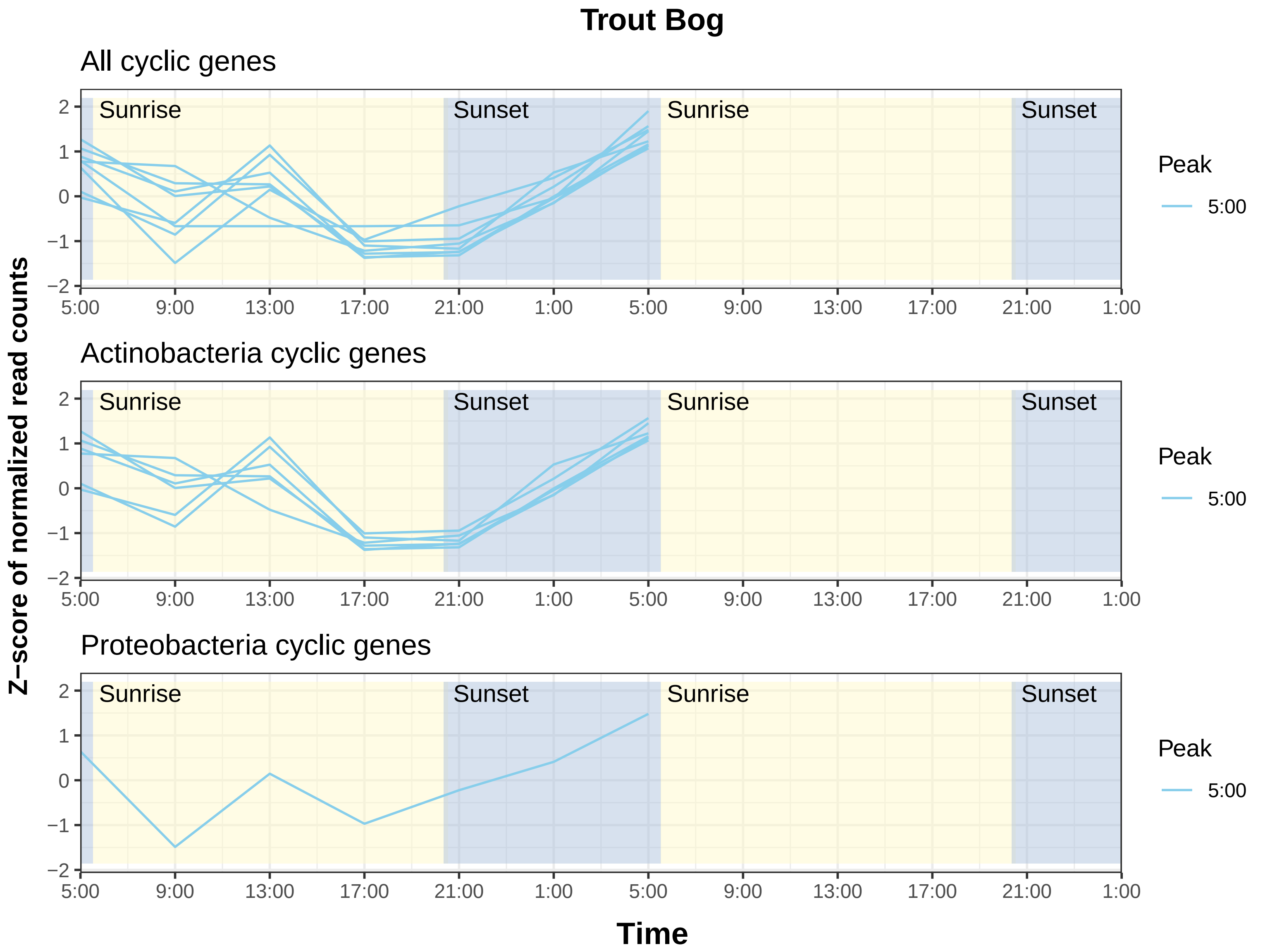

Trout Bog
All cyclic genes
Actinobacteria cyclic genes
Proteobacteria cyclic genes
Sunrise
Sunset
Sunrise
Sunset
Peak
5:00
Z−score of normalized read counts
Time
5:00
9:00
13:00
17:00
21:00
1:00
5:00
9:00
13:00
17:00
21:00
1:00
2
1
0
−1
−2
