## Supplemental Figures 2 to 5 for "Diversity, distribution, and expression of opsin genes in freshwater lakes"

**This file only has Figures S2, S3, S4 and S5. Figure S1 is a separate PDF file with its legend shown below.**

**Figure S1.** A rectangular view of the phylogenetic tree shown in Figure 1. Bootstrap values were calculated based on 100 replicates. Reference sequences from the public databases are labeled in black, with their taxonomic classification indicated and IMG gene IDs or GenBank accession numbers listed in the bracket. Opsin genes from our metagenomes were labeled in blue for ME and red for TE and TH. Sequences in MAGs/bins were highlighted with bold fonts, with their taxonomic classification indicated and IMG gene IDs listed in the bracket. For the rest (un-binned) metagenome sequences, gene names are their sequence Locus ID in IMG. Locus ID starting with "B570J40625", "TBL_comb48_EPIDRAFT", and "TBL_comb47_HYPODRAFT" are from ME, TE and TH metagenomes, respectively.

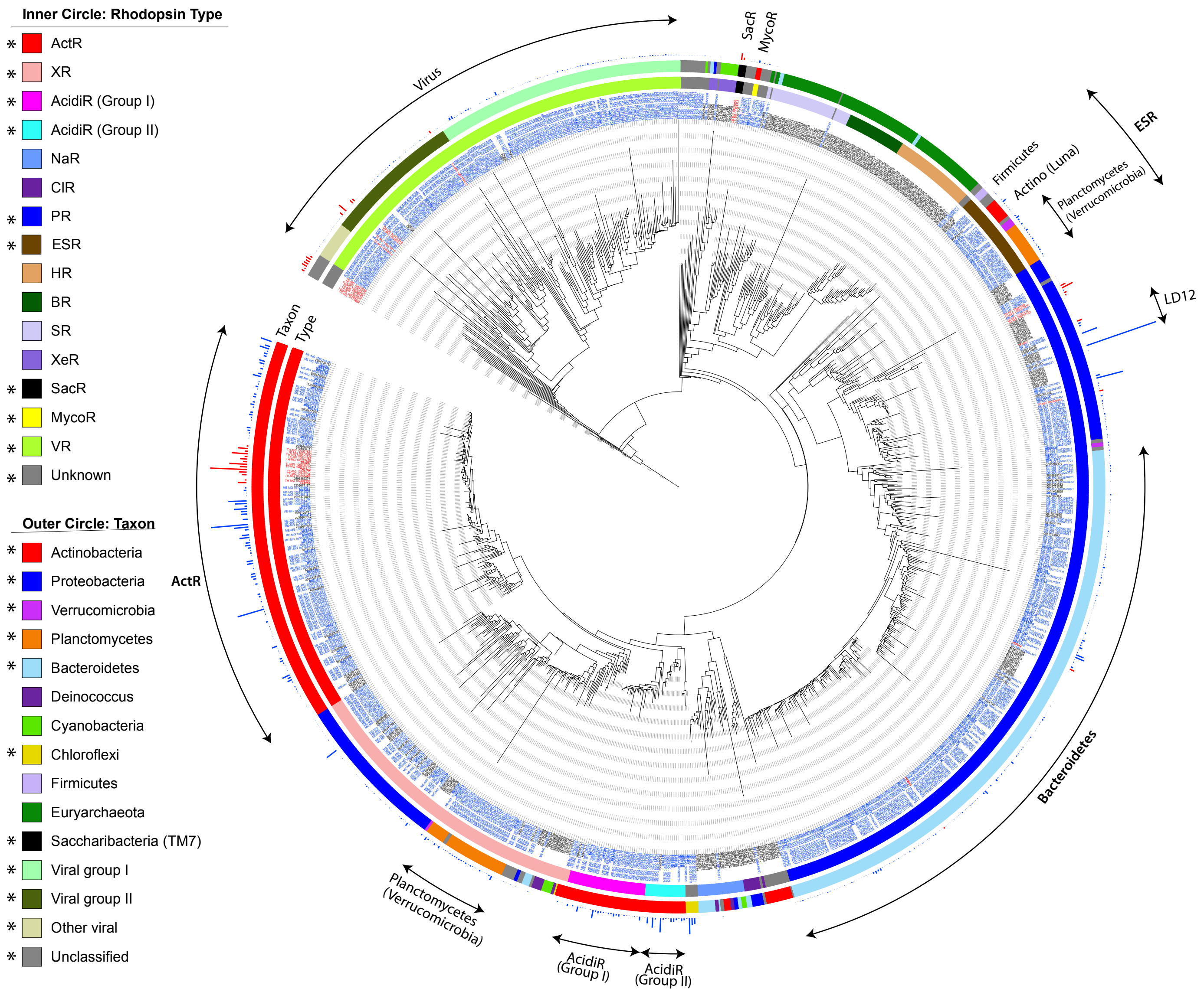


**Figure S2.** Summary phylogenetic tree with metagenome sequences shorter than 200 amino acids added to the phylogenetic tree shown in Figure 1 by using PPlacer. Opsin gene names are colored as following: blue for ME, red for TE and TH metagenomes, and black for references from the database. For sequences in MAGs/bins (highlighted with bold fonts), gene names start with "ME", "MEE", "TE", and "TH" respectively. For the rest (un-binned) metagenome sequences, gene names start with "ME-MG", "TE-MG", and "TH-MG", respectively. The inner circle color strip indicates rhodopsin types, and the outer circle color strip indicates taxon. Rhodopsin types and taxon present in our metagenomes were indicated with "*" in the legend. The outmost bar indicates normalized coverage-weighted abundance of opsin genes in the metagenome (blue for ME, and red for TE and TH). Rhodopsin types include ActR (actinorhodopsin), XR (xanthorhodopsin), AcidiR (acidirhodopsin), NaR (sodium-pumping rhodopsins), ClR (chloride-pumping rhodopsins), PR (proteorhodopsin), ESR (*Exiguobacterium sibiricum* rhodopsin), HR (halorhodopsin), BR (bacteriorhodopsin), SR (sensory rhodopsin), XeR (Xenorhodopsin), SacR (Saccharibacteria rhodopsin), MycoR (DTG rhodopsins represented by the freshwater Myco tribe), and VR (viral rhodopsin).

**Figure S3.** Percent distribution of opsin genes among different rhodopsin types and different taxonomy within the three combined assemblies. As a fraction of opsin genes in the metagenomes are incomplete, for opsin genes shorter than 200 amino acids (i.e. ~80% of the microbial rhodopsin hidden Markov model, HMM, length), their coverage-weighted abundance was further normalized using their recovered lengths relative to the bacteriorhodopsin HMM length so that the abundance of these gene fragments can be compared to complete and near-complete opsin genes to estimate their percent distribution.

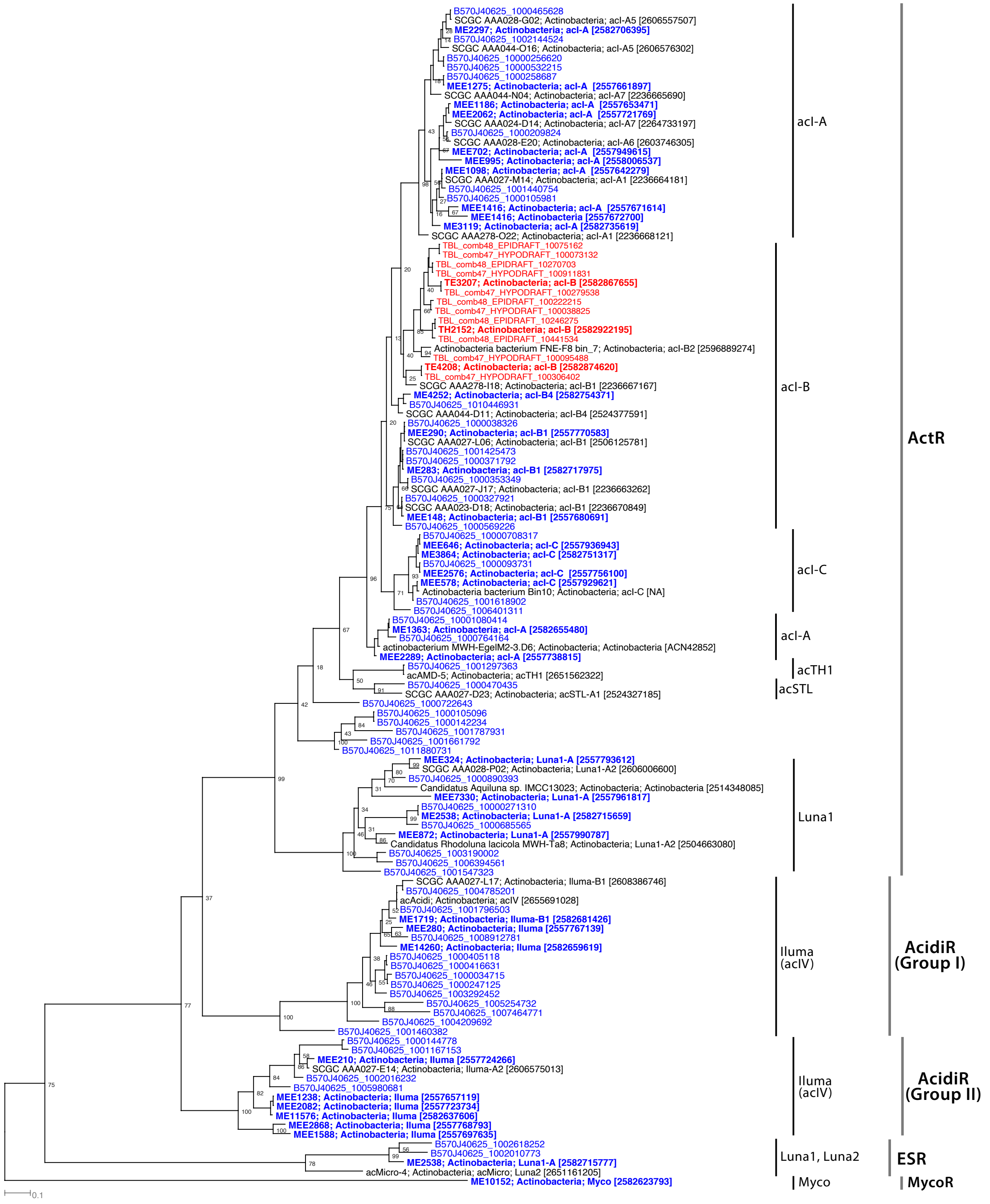


**Figure S4.** Subtree of opsin sequences belonging to freshwater Actinobacteria. Opsin genes from our metagenomes were colored (blue for ME and red for TE and TH). Sequences in MAGs/bins from our study were highlighted with bold fonts, with their taxonomic classification indicated and their opsin sequence IMG gene IDs listed in the bracket. For the remaining metagenome sequences (un-binned), gene names are their sequence Locus ID in IMG. Locus ID starting with "B570J40625", "TBL_comb48_EPIDRAFT", and "TBL_comb47_HYPODRAFT" are from ME, TE and TH metagenomes, respectively.

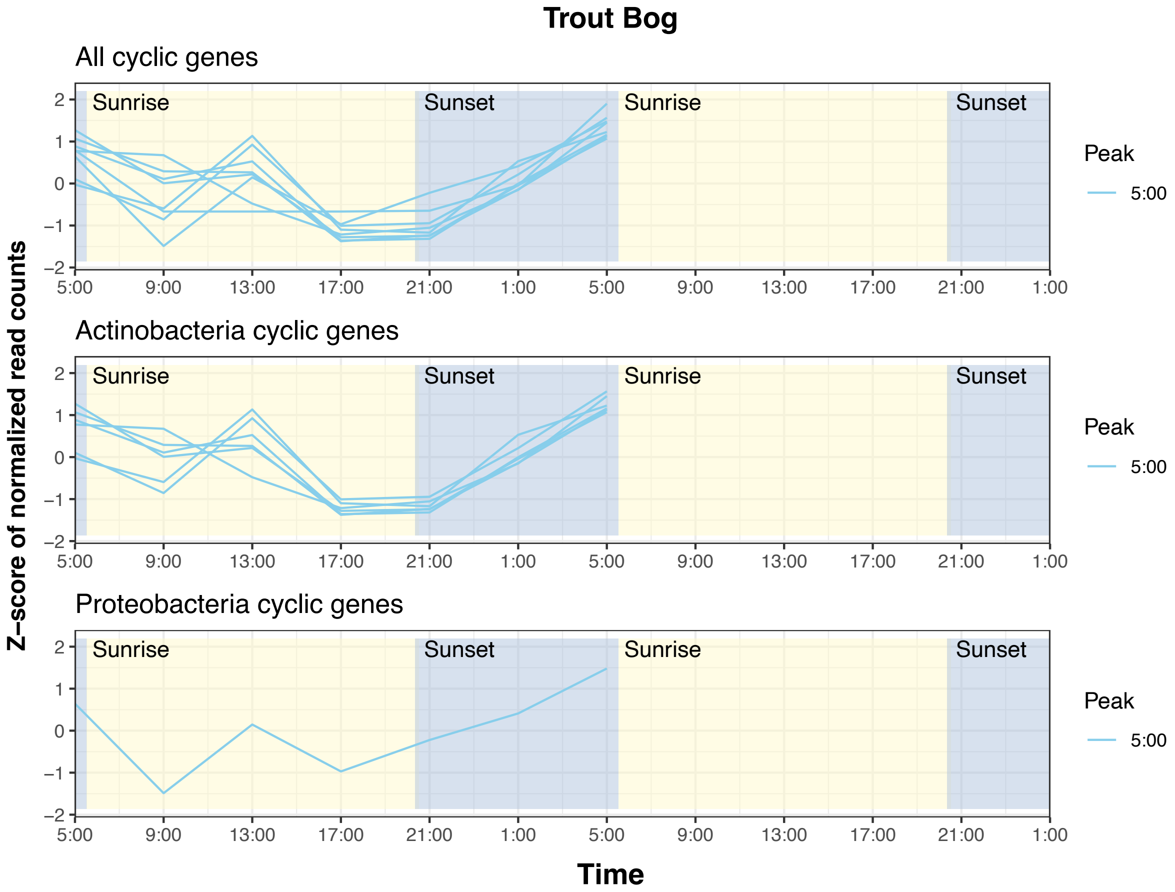


**Figure S5.** Diel cyclic trends of opsin gene expression. Cyclic genes were determined based on expression data collected from one day period using the RAIN test with the *p*-value of < 0.05. The top panel shows all cyclic genes, and the bottom three panels show cyclic genes in Actinobacteria and Proteobacteria. Read counts in transcripts per liter were z-score transformed and are shown on y-axis. Genes were colored by the peak hour of their expression as determined in the RAIN test, with warm and cold colors for peaks at "day" and "night" (as determined by photosynthetically active radiation), respectively.
