## Supplemental Text for "Diversity, distribution, and expression of opsin genes in freshwater lakes"

**Supplementary Text – Viral origin of MEE3901, MEE6525 and MEE15704 bins**

Among the 92 opsin gene-containing MAGs/bins, MEE3901, MEE6525 and MEE15704 were very likely derived from viruses. The three bins have sizes ranging from 0.43 to 0.62 Mbp and contain almost no bacterial conserved single-copy genes (CSCGs). Their genome statistics from on the IMG annotation pipeline (such as the low GC content, the extremely low percentages of genes with COG, pfam, TIGRfam, and KO annotation, and the absence of bacterial CSCGs) are much more similar to viral genomes, such as *Phaeocystis globosa* viruses, as compared to the 89 bacterial MAGs/bins included in this study (**Table S8**). In addition, many genes in the three bins (ranging from 34% to 50% of all protein-coding genes in the bin) are homologous to *Phaeocystis globosa* viruses (**Table S8**), suggesting that these three bins are likely of viral origin.

We therefore conducted hmmsearch (HMMer v.3.2.2) to search for virus orthologous groups (VOG) (http://vogdb.org) in the three bins. In each of the three bins, we identified phage hallmark genes whose functions are critical for phage morphology and bacterial infection. These genes include capsid protein, tail proteins (spike, fiber, collar, assembly, etc.), baseplate proteins, virion membrane proteins, viral transcription factors, *etc*. (**Table S9**). The presence of these phage hallmark genes in the three bins supports their viral origin.
